# Enhancing the Reverse Transcriptase Function in Taq Polymerase via AI-driven Multiparametric Rational Design

**DOI:** 10.1101/2024.07.24.604875

**Authors:** Yulia E. Tomilova, Nikolay E. Russkikh, Igor M. Yi, Elizaveta V. Shaburova, Viktor N. Tomilov, Galina B. Pyrinova, Svetlana O. Brezhneva, Olga S. Tikhonyuk, Nadezhda S. Gololobova, Dmitriy V. Popichenko, Maxim O. Arkhipov, Leonid O. Bryzgalov, Evgeny V. Brenner, Anastasia A. Artyukh, Dmitry N. Shtokalo, Denis V. Antonets, Mikhail K. Ivanov

## Abstract

Modification of natural enzymes to introduce new properties and enhance existing ones is a central challenge in bioengineering. This study is focused on the development of Taq polymerase mutants that show enhanced reverse transcriptase (RTase) activity while retaining other desirable properties such as fidelity, 5′-3′ exonuclease activity, effective deoxyuracil incorporation, and tolerance to locked nucleic acid (LNA)-containing substrates. Our objective was to use AI-driven rational design combined with multiparametric wet-lab analysis to identify and validate Taq polymerase mutants with an optimal combination of these properties. The experimental procedure was conducted in several stages: 1) On the basis of a foundational paper, we selected 18 candidate mutations known to affect RTase activity across six sites. These candidates, along with the wild type, were assessed in the wet lab for multiple properties to establish an initial training dataset. 2) A ridge regression model was trained on this dataset to predict the enzymes’ properties. This model enabled us to select 14 new candidates for further experimental testing. 3) We refined our predictive model using Gaussian process regression and trained it on an expanded dataset now including 33 data points. 4) Leveraging the refined model, we screened *in silico* over 27 million potential mutations, thus selecting 16 for detailed wet-lab evaluation. Through this iterative data-driven approach, we identified 18 enzymes that not only manifested considerably enhanced RTase activity but also retained a balance of other required properties. These enhancements were generally accompanied by lower K_d_, moderately reduced fidelity, and greater tolerance to noncanonical substrates, thereby illustrating a strong interdependence among these traits. Several enzymes validated via this procedure were effective in single-enzyme real-time reverse-transcription PCR setups, implying their utility for the development of new tools for real-time reverse-transcription PCR technologies, such as pathogen RNA detection and gene expression analysis. This study illustrates how AI can be effectively integrated with experimental bioengineering to enhance enzyme functionality systematically. Our approach offers a robust framework for designing enzyme mutants tailored to specific biotechnological applications. The results of our biological activity predictions for mutated Taq polymerases can be accessed at https://huggingface.co/datasets/nerusskikh/taqpol_insilico_dms.

## Introduction

Modification of natural enzymes in order to give them new properties and to enhance existing ones is a rapidly developing field of bioengineering [1] [2] [3] [4]. Demands of modern science, industry, and health care stimulate the search for new enzymes or improvements of known enzymes to implement new or better features useful in these fields. Early on, random mutagenesis followed by selection that mimics natural evolution was the most widespread way to obtain new functional mutants of enzymes (proteins with modified functions). This approach, while being effective enough, is time-consuming and labor-intensive. Later, rational-design approaches began to emerge, based on information obtained from protein alignments and structures and structures of their complexes with ligands. Although traditional rational design has substantially advanced enzyme engineering, it is often based on extensive empirical data and can be limited by the complexity of protein interactions. As the quest for efficiency and precision in protein engineering continues, the integration of computational tools has become inevitable.

Deep learning, especially through the use of language models, has transformed protein science by enabling researchers to efficiently harness vast genomic databases like BFD and Uniref50. These protein language models (PLMs), trained via unsupervised pretraining techniques, predict masked amino acids by interpreting contextual information from visible sequence data. The resulting embeddings — dense, information-rich vectors for each amino acid — capture essential biophysical and structural properties not explicitly mentioned in the data. Through aggregation of these embeddings, entire protein sequences can be parameterized, laying a comprehensive foundation for predicting mutations’ effects on a protein’s structure. Such capabilities allow for practical application of these models in various bioengineering tasks, including prediction of secondary structures, of residue contacts, and of a mutational impact on enzyme functionality [5] [6] [7] [8] [9] [10], thus advancing the field beyond previously available methods.

The first remarkable application of a PLM to rational enzyme engineering was achieved with UniRep [11], which was trained on over 20 million protein sequences from UniRef50, thereby allowing the model to learn general protein features in an unsupervised manner. The representations learned by UniRep effectively support linear regression models in guiding directed evolution, thus proving adequate for capturing necessary information about mutant proteins [11]. It has also been found that UniRep simulates a fitness landscape accurately enough for engineering applications using only 24 functionally characterized proteins bearing amino acid substitutions [12]. UniRep utilizes a deep recurrent neural network based on the biLSTM architecture, which is relatively small by modern standards. In contrast, current large language models, especially those in protein science, predominantly involve the transformer architecture, which has been demonstrated to be superior in handling complex sequence data. Among modern families of PLMs, we can mention ESM [13] [14], RITA [15], ProtT5 [5], ProstT5 [16], ProGen2 [17], ProtGPT2 [18], and ECNet [19], representing the current state of the art in PLMs and yielding promising results in various applications including protein design. Alongside our primary use of PLMs for predicting protein behaviors, multiple sequence alignment (MSA)-based models, also known as MSA transformers [9] [10], introduce a unique approach to integrating evolutionary information during protein analysis. These models require constructing MSAs as input, thereby effectively taking two-dimensional (2D) input instead of traditional 1D data. This 2D input necessitates a specialized architecture, significantly increasing computational and memory demands.

Our study was intended to harness these advanced modeling techniques in order to predict and experimentally validate new enzyme mutants having a combination of essential properties paving the way for more effective biotechnological tools. We believe that the most efficient approach to rational protein design is an iterative procedure involving several rounds of computational prediction and subsequent experimental studies, especially when tradeoffs must be made between several simultaneously required biological functions and physicochemical characteristics because designing new proteins is often a complex problem with many optimization criteria. To prove this concept, we attempted to construct and experimentally validate a predictive model aimed at selecting mutants of *Thermus aquaticus* DNA polymerase (hereinafter referred to as Taq pol)—having a set of required properties—by combining structure-based rational design and experimental results of multiparametric wet-lab assays.

Thermostable DNA-dependent DNA polymerases are among the most popular objects of protein engineering because they are a key component of nucleic acid amplification techniques and are widely used as modern tools in molecular biology and biotechnology. Out of these enzymes, Taq pol has found widespread applications due to a combination of numerous useful properties, such as ease of accumulation in bacterial expression systems, extreme thermostability, and strong 5′-3′ exonuclease activity. At the same time, Taq pol is characterized by relatively low fidelity (because it lacks an active 3′-5′ exonuclease proofreading domain) and negligible strand displacement activity and has only very low intrinsic RNA-dependent DNA polymerase activity. This limits the use of wild-type (WT) Taq pol as a core enzyme in a number of molecular biological applications. At the same time, Family A DNA polymerases, including Taq pol, are known for their structural plasticity: they tolerate multiple amino acid substitutions, even in evolutionarily conserved regions [20]. This property makes Taq pol a convenient model for function enhancement and modification studies. Numerous modified versions of Taq pol have been obtained that are characterized for example by increased resistance to PCR inhibitors [21], improved elongation [22] and strand displacement abilities [23] [24], 3′-5′ exonuclease activity [25], a reduced capacity to discriminate against dideoxynucleotides [26] or to elongate mismatched PCR primers [27][28], wider substrate specificity [29] [30], or cold sensitivity [31].

One of research fields in Taq pol engineering is the creation of enzymes with dual DNA- and RNA-dependent DNA-polymerase (reverse transcriptase, RTase) activities for biotechnology, molecular genetic studies, and applied diagnostic tools. Traditional mesophilic RTases such as M-MuLV or AMV have difficulty synthesizing cDNA through stable RNA secondary structures and GC-rich motifs. High-temperature reverse transcription (RT) may enhance cDNA synthesis efficiency and reduce primer dimerization and side product formation, thus making primers more sequence-specific. RTases derived from Taq pol, known also for its inhibition tolerance, are expected to retain these properties and thus to have an advantage over mesophilic RTases. The single-enzyme approach could streamline the development of real-time RT-qPCR tools, also benefiting from the enzyme’s thermostability in terms of storage, shipping, and automation. These and other potential benefits stimulate the ongoing efforts to create new high-temperature RTases engineered from Taq pol and other thermostable DNA polymerases. Successful examples of enhancement of the RTase activity in Taq pol via introduction of one or more point mutations have long been known [32] [33] [34] [35] [36] [37] [24] [38]. Of note, in different articles, modifications at completely different sites have had similar effects on this property. For instance, in ref. [34], a directed-evolution experiment with the Stoffel fragment of Taq pol gave 27 mutant enzymes showing appreciably (up to two orders of magnitude) enhanced RTase activity, which was achieved for different mutants through amino acid substitutions at more than 50 positions. In the majority of these projects, enzymes with enhanced RTase activity have been selected based on a limited set of parameters from a huge number of mutants obtained by randomized mutagenesis. The latter typically generates a large pool of candidate enzymes, which has to be investigated in detail to choose the most promising mutants.

Given that multiparametric evaluation of an original pool of candidates by wet-lab experiments is time-consuming and labor-intensive, initial sorting of candidates is traditionally done in a limited series of experiments. As a consequence, possible negative effects of the introduced mutations on some important properties of the modified enzyme can be overlooked, and, on the other hand, the main improved characteristic can be evaluated under suboptimal conditions. For this reason, some promising candidates can be rejected by the initial selection, and conversely, candidates with undesired properties can pass through the selection screen because the same amino acid substitution at a functionally significant position can simultaneously affect several functions, sometimes in opposite directions. In this regard, in our opinion, an alternative strategy looks attractive: *in silico* preselection of candidate mutants by means of rational predictions about an extended range of biotechnologically significant properties, to take them into account right from the start. Accordingly, we set the following selection goal: to create an enzyme that a) possesses an enhanced RTase activity in a PCR-compatible buffer (obviating manganese ions), and b) retains some beneficial properties of WT Taq pol (sufficient fidelity, 5′-3′ exonuclease activity, thermal stability, the ability to effectively incorporate dU, and the capacity to process locked nucleic acid (LNA)-containing substrates; hereafter: LNA substrates) that are unaffected or even improved. A list of parameters was compiled based on the suitability of mutants for molecular diagnostic applications. Additionally, we tested all the produced mutant enzymes for the “hot start” capacity through blocking by a monoclonal antibody or a DNA aptamer.

Given the impracticality of constructing MSAs for the millions of candidates we evaluated, direct applications of PLMs were chosen that efficiently integrate evolutionary insights without the extensive resource burden associated with MSA transformers. By means of several rounds of experiments, we created a set of multivariate regression models built on top of embeddings obtained via the ProtT5 language model family [5]. This PLM-based regression model allowed us to identify several Taq pol mutants with noticeably enhanced RTase activity combined with the preservation (or even enhancement) of a number of required preselected characteristics.

## Methods

### Starting the selection of substitution mutants

To train the first version of the model aimed at predicting effects of amino acid changes on the selected Taq pol parameters, we began with a multiparametric wet-lab testing of a limited set of Taq pol mutants carrying various single amino acid substitutions that were located at key positions in different structural domains and were expected to affect various protein properties in different directions. Initially, we applied the following restrictions: for resource-saving reasons, to characterize only 18 mutants first, together with the WT protein serving as a reference; to check several key positions localized to all structural domains (thumb, palm, and finger); and to include a few mutants that have low expected activity in PCR. Thus, using a limited number of options, we intended to obtain various combinations of substitutions in terms of changes in such physicochemical properties as charge loss/replacement, physical volume, aromaticity, and polarity.

The results from [37] were employed as a starting point for choosing the substitutions to be evaluated experimentally. There, 12 key positions were investigated in detail: N483, E507, S515, and K540 (thumb); A570, D578, V586, and V783 (palm); and I614, H639, F667, and M747 (finger). We selected six out of these 12 positions: M747 and D578 (contacting the template strand), E507 and S515 (coming into contact with the primer strand), and H639 and F667 (binding to an incoming nucleoside-5′-O-triphosphate). Other positions were excluded due to their high tolerance to amino acid substitutions. Thus, the following 18 mutations were finally chosen for the first round of wet-lab assays: E507D, E507Q, E507K, S515D, S515F, S515N, D578K, D578N, D578W, H639A, H639F, H639Y, F667A, F667Y, F667M, M747E, M747D, and M747Q. Six conservative substitutions (E507D, E507Q, S515D, D578N, H639Y, and F667Y) and four substitutions expected to result in PCR activity loss (S515F, H639A, F667A, and M747D) according to ref. [37] were intentionally included in this list. Locations of the investigated amino acid positions in 3D structure of the Taq pol large fragment complexed with a DNA molecule are depicted in **Figure 1** (together with other positions further assayed in our study).

**Figure 1.**
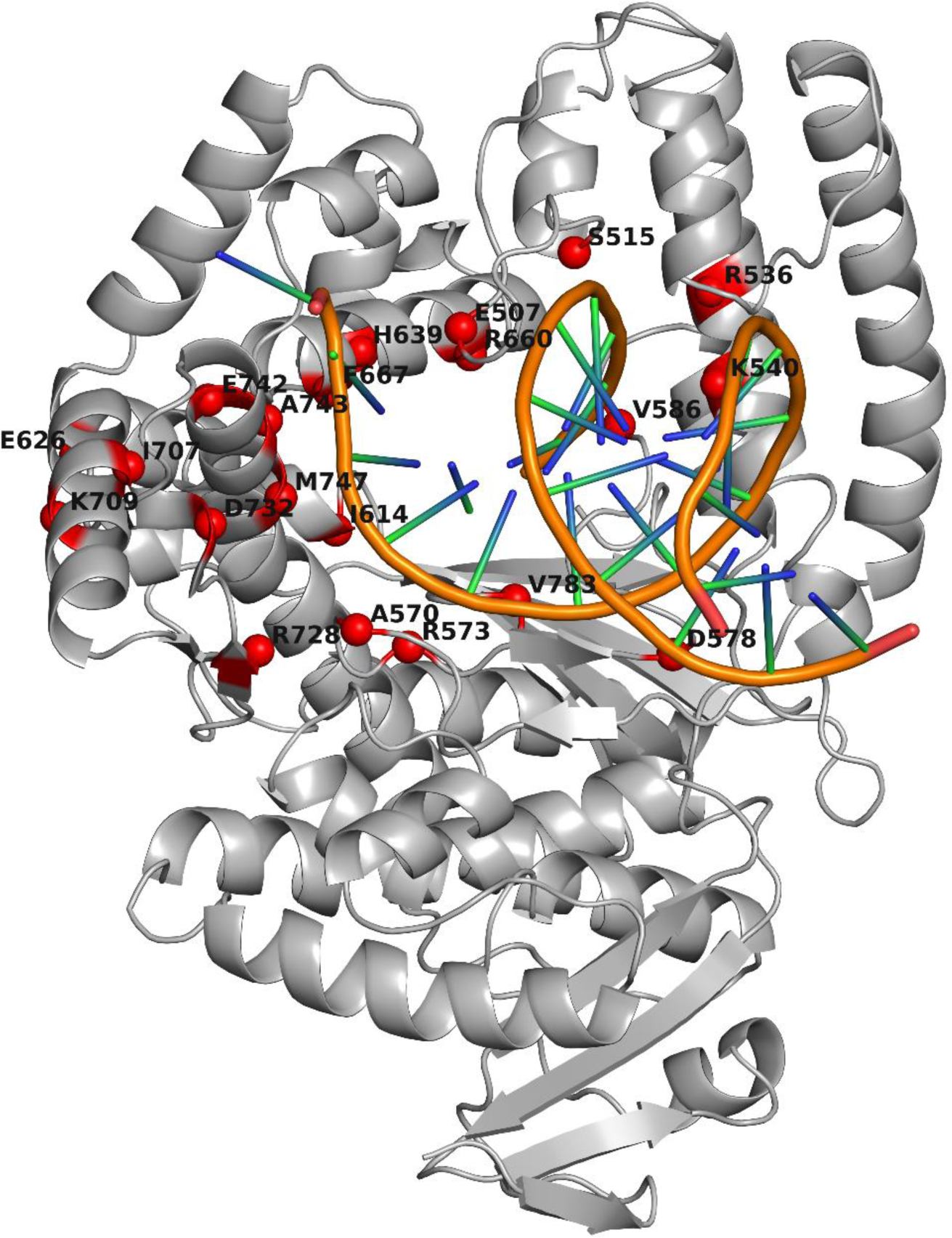
Locations of assayed amino acid positions in 3D structure of the large fragment of *T. aquaticus* DNA polymerase I complexed with a DNA molecule (Protein Data Bank ID: *3KTQ*). The spatial structure of Taq pol is presented as a gray ribbon diagram. Red spheres denote Cα atoms of amino acid residues (aa) that were mutated alone or in several combinations. The labels indicate residues and their positions in WT Taq pol (SwissProt: *P19821*). The image was produced in PyMOL v.2.5.0 [39].

### Generation of Taq pol mutants

The nucleotide sequence of the Taq pol–encoding gene was codon-optimized for translation in the *Escherichia coli* expression system. All rare codons were eliminated, and codon frequency after the optimization was equal to or greater than 8/1000. The coding part of the synthetic gene sequence started from the fourth codon of the original Taq pol sequence: therefore, the protein we used differed from native Taq pol by the absence of the first 3 aa. Nonetheless, to facilitate the comprehension and interpretation of the results in the context of publications by other research groups, the numbering of amino acid positions corresponding to native Taq pol is utilized throughout the text. Oligonucleotides for PCR-based gene synthesis were designed in online software DNAWorks (v3.2.4) (https://hpcwebapps.cit.nih.gov/dnaworks/). The gene sequence was divided into five fragments ∼500 bp long and flanked with restriction endonuclease sites, which helped to clone the gene into the pJET1.2/blunt vector (Thermo Fisher Scientific, USA). The mutations were introduced into the corresponding fragments by site-directed mutagenesis. The full-length gene for each Taq pol mutant was assembled by sequential ligation of fragments into the pET28a-Novagen expression vector (Sigma-Aldrich, USA). All sequences of synthetic constructs were verified by Sanger sequencing on an Applied Biosystems 3500 instrument (Thermo Fisher Scientific) with the BigDye Terminator v3.1 Cycle Sequencing Kit. All Taq pol mutants were expressed in *E. coli* BL21(DE3)pLysS. Competent cells were transformed with the expression plasmid and grown at 37 °C in 400 ml of the Luria–Bertani (LB) medium containing 30 μg/ml kanamycin and 34 μg/ml chloramphenicol. When cell density reached OD_600_ of 0.8–0.9, protein expression was induced with 1 mM isopropyl β-D-1-thiogalactopyranoside. After 3 h of expression, the cells were centrifuged at 6400 g for 10 min at 4 °C and stored at −70 °C until analysis. Cell pellets were resuspended in a buffer consisting of 20 mM NaH_2_PO_4_ (pH 8.0), 50 mM NaCl, 0.1 mM EDTA, 1 mM PMSF, and 1% of Triton X-100. After heat denaturation at 75 °C for 40 min in a water bath, the lysates were centrifuged at 39,000 × *g* and 4 °C for 20 min. The supernatant was treated with 0.05% polyethyleneimine to remove chromosomal DNA and centrifuged at 39,000 × *g* and 4 °C for 20 min. The supernatant was passed through a 0.45 μm syringe filter and purified on a Ni-NTA column using a buffer composed of 20 mM NaH_2_PO_4_ (pH 8.0), 0.5 M NaCl, 0.1 mM EDTA, 15 mM imidazole, and 0.1% of Triton X-100. The proteins were eluted with elution buffer (20 mM NaH_2_PO_4_ pH 8.0, 0.5 M NaCl, 0.1 mM EDTA, 250 mM imidazole, and 0.1% of Triton X-100) and then dialyzed against another buffer (20 mM Tris HCl pH 8.0, 75 mM NaCl, and 0.1 mM EDTA) overnight with stirring in a cold room. The concentration of purified proteins was determined spectrophotometrically.

### Agarose gels

Horizontal electrophoresis chamber Wide Mini Sub Cell GT and Gel Documentation System Gel Doc XR+ (Bio-Rad, USA) were used to examine 2% agarose gels during electrophoresis and to take photographs.

### Oligonucleotide primers and probes

The oligonucleotides were designed using the PrimerQuest online service (https://eu.idtdna.com/calc/analyzer). All oligonucleotides were synthesized at AO Vector-Best. Structures of primers and fluorescently labeled probes are presented in Table **S1**.

**Kinetic curves of the change in fluorescence anisotropy** in kinetic assays of the formation of enzyme–substrate complexes (determination of equilibrium dissociation constants: K_d_) as well as kinetic curves of the change in SYTO 13 fluorescence in experiments on the rate of polymerase-driven synthesis (determination of catalytic constants of the polymerization reaction rate in the presence of dT and dU: k_Cat_(dT) and k_Cat_(dU)) were recorded on an SX20 stopped-flow spectrometer (Applied Photophysics, UK) with the Pro-Data SX20 software (Applied Photophysics). Each kinetic curve was obtained by averaging 15 to 30 experimental curves. The fluorescence anisotropy measured in the experiment at each time point depends on the current concentration of the complex of polymerase with a labeled substrate called H-Kd (see **Table S1**): the higher the concentration of the specified complex, the higher the anisotropy value is. Quantitative processing of the experimental data was carried out using the “minimize” function of the scipy.optimize Python library via optimization of the parameters determining the formation of the “polymerase-labeled oligonucleotide” complex (in the case of K_d_ calculation) and through optimization of the parameters determining the elongation rate of the 20_60 hairpin template (in the case of k_CatT_ and k_CatU_ calculation). Reaction conditions were as follows: cell volume, 20 µl; temperature, 55 °C; buffer: 5 mM Tricine-KOH pH 8.0, 100 mM KCl, 3.4 mM MgCl2, 0.1 mM each dNTP, 0.01% of Tween 20, 1/5000 SYTO 13, and 20 nM hairpin template. Fluorescence was detected at an excitation wavelength of 535 nm and a cutoff filter >570 nm (in the case of K_d_ determination) or using an excitation wavelength of 485 nm and a cutoff filter >530 nm (in the case of k_CatT_ and k_CatU_ determination).

### Assessment of the hot-start capacity by means of a monoclonal antibody and DNA aptamer

This experiment with a monoclonal antibody to Taq pol (Clontech, USA) and a DNA aptamer (Vector-Best, Russia) was performed as described elsewhere [40]. Briefly, polymerase kinetics and their change related to the blocking by the antibody or aptamer were registered by means of an increase in SYTO 13 fluorescence after elongation of the OnOff hairpin template (see **Table S1**) during catalysis by different concentrations of tested enzymes at different temperatures. Hereinafter (if not specified otherwise), in all kinetic assays, a 1/2500 dilution of SYTO 13 from a commercial stock solution (Thermo Fisher Scientific) was used. The reaction was carried out in 1× RT buffer as 500 cycles at 45 °C. Blocking was considered effective when it resulted in at least a 10-fold decrease in the reaction rate.

**The effective reaction rate constant** was evaluated under the same conditions as in the hot-start experiment, via 500 cycles at 60 °C. For each enzyme, four concentrations were analyzed in serial 2-fold dilutions. Prior to this, the lowest concentration of each enzyme was experimentally found that gave a kinetic curve of the classic shape. Each enzyme concentration was analyzed in triplicate, and the experiment with each enzyme was conducted twice: thus, for each tested concentration of each enzyme, six kinetic curves were built. The inverse kinetics problem was examined for each curve dataset via calculation of the effective reaction rate constant. The inverse problem was solved by a numerical approach involving Gauss and RKM methods for a system of differential equations and “the Peak descent” method for step-by-step curve approximation [41]. The effective rate constants obtained in this way were averaged for each enzyme. Finally, the ratio of the activity of each mutant enzyme to the control was computed:

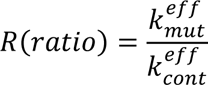

**The influence of dTTP substitution with dUTP on the synthesis rate of the DNA-dependent DNA polymerase** was evaluated under the same conditions. The second and third concentrations of each enzyme from the serial dilutions described above were tested. Each enzyme concentration was analyzed in triplicate. The experiment with each enzyme was conducted in two ways: either in 1× RT buffer or in its modification where dTTP was replaced with dUTP at the same concentration. Data were recorded as the ratio of effective reaction rate constants between reactions in dTTP-containing buffer and dUTP-containing buffer (hereinafter: the dT/dU rate).

**RTase activity** was assessed in two ways:

1. Through direct evaluation of RT kinetics by means of the increase in SYTO 13 fluorescence after elongation of a double-stranded substrate comprising 7.5 nM oligo(dT) primer and 600 pg/μl poly(r)A in RT buffer. The 50 μl reaction was carried out in 1× RT buffer (50 mM tricine-KOH pH 8, 50 mM KCl, 0.0001% of Tween 20, 14.5% of trehalose, 100 μg/ml BSA, 240 μM dTTPs, and 3 mM MgCl_2_) on a CFX96 thermocycler with an optical module (Bio-Rad) by the following protocol: 500 cycles at 45 °C. Enzyme concentrations were selected so that the kinetic curve could be clearly recorded and had the classic shape. Each experiment was conducted in duplicate, and curves plotted from the mean values for each experimental data point were employed for interpretation. Ratios of the activity of each mutant enzyme to controls (which were the WT and subunit p66 of HIV RTase, hereinafter referred to as p66) (AO Vector-Best, Russia) were computed as described above.
2. Via evaluation of cDNA synthesis efficiency of different Taq pol mutants (including the WT) on specific RNA targets of various lengths. The RT (30 min at 60 °C) was allowed to proceed in a 50 μl reaction mixture composed of 1× PCR buffer (Vector-Best, Russia) supplemented with 10% of trehalose, 100 μg/ml BSA, 5 mM MgCl_2_, 0.4 mM each dNTP, 0.6 M betaine, and 500 nM corresponding sequence-specific reverse primer (see **Table S1**). Synthetic RNA transcripts containing specific sequences of the human parainfluenza virus 2 (HPIV2) phosphoprotein gene (90 nt), human metapneumovirus (hMpV) nucleoprotein gene (116 nt), rhinovirus (RhV) 5′-UTR (201 nt), and human immunodeficiency virus (HIV-1) 5′ LTR (526 nt) were added at the same concentration (∼10^6^ copies/reaction) to spike the reaction mixture. p66, serving as the reference enzyme, was assayed under the same conditions, except for the temperature and enzyme concentration, which were selected in preliminary experiments (15 nM and 50 °C). Five microliters of each RT reaction were mixed with respective sets of primers and fluorescently labeled probes (See **Table S1**) in nucleic acid elution buffer (Vector-Best) added to the final volume of 50 μl, and then amplification was performed by real-time PCR with the help of ready-to-use freeze-dried master mixes containing PCR buffer and WT Taq pol (Vector-Best). The thermal cycling program was as follows: 2 min at 95 °C and then 50 cycles of 10 s at 94 °C and 20 s at 60 °C. Amplification data were recorded on the CFX96 instrument. Each experiment was conducted twice, and the obtained C_q_ values were averaged. RTase activity in each “Taq pol mutant/cDNA” pair was assessed as the difference between the average C_q_ value shown by the tested enzyme and the value shown by the p66 RTase.

**Evaluation of the ability to process LNA substrates** was performed under the same conditions. The second concentration for each enzyme from the serial dilutions described above was utilized. Each enzyme was tested in triplicate with each of four hairpin substrates: LNA-0, LNA-1, LNA-2, and LNA-3 (all of which had identical structure, but LNA-1–3 carried LNA nucleotides at different positions, see **Table S1** and **Fig. S1**). The kinetic curves were averaged for each enzyme/substrate pair. The change in the reaction rate with each modification (LNA nucleotide) in the substrate for each mutant enzyme was normalized to the change in the reaction rate with the same modification for the WT enzyme.

### Assessing the usability of selected Taq pol mutants in a single-tube RT-PCR with TaqMan detection

Enzymes with the best combination of desirable characteristics (sufficient DNA-dependent DNA polymerization rate and PCR efficiency, enhanced RTase activity, the ability to utilize LNA substrates and dUTP, and the capacity to effectively cleave TaqMan probes) were tested for suitability for single-tube RT-PCR. The 50 μl reaction was carried out on the CFX96 thermocycler in 1× PCR buffer with primers to the target sequences at a concentration of 500 nM each and fluorescently labeled probes at 250 nM. Each mutant enzyme was added at 120 nM, and synthetic RNA transcripts carrying the 90-nt HPIV2 sequence and the 116-nt hMpV sequence were used to spike the reaction mixture at 10^6^ and 10^4^ copies/reaction. Either a mixture of 26 nM p66 and 120 nM WT or 120 nM WT alone served as a control. The following thermal cycling conditions were implemented: 15 min at 60 °C (found for all mutant Taq pol mutants in preliminary optimization experiments) or 15 min at 50 °C (for p66), then 1 min at 94 °C and 50 cycles of 10 s at 94 °C and 20 s at 60 °C with fluorescence recording. The data were interpreted with respect to two parameters: C_t_ reflected integral efficiency of RT and cDNA amplification, and the amplitude and shape of the fluorescence curve signified the 5′ exonuclease activity resulting in the cleavage of the labeled probe.

**Fidelity of DNA-dependent DNA polymerase activity of the mutant polymerases** was evaluated as the proportion of misincorporated nucleotides during synthesis of a specific 99 bp fragment of the hepatitis B virus (HBV) *S* gene, as assessed by examination of next-generation sequencing data. Purified HBV DNA (7.5 pg) served as a template. HBV genome fragments were synthesized on Bio-Rad CFX96 in a 30 μl reaction consisting of 1× PCR buffer with primers HBV-F and HBV-RR at 500 nM each and the fluorescently labeled HBV-P probe at 250 nM according to the following program: 2 min at 95 °C, 10 s at 94 °C, and 20 s at 60 °C. Taq pol mutants were added at 120 nM. The reaction was stopped at the time of reaching a plateau (for different enzymes, it corresponded to different numbers of amplification cycles). Polymerase-driven synthesis was monitored using SYTO 13 fluorescence (to track the reaching of a plateau) and ROX fluorescence (to assess PCR efficiency and the ability to cleave 5′-labeled probes). It is noteworthy that the PCR efficiency determined in this analysis was applied as an independent criterion to model training and selection of candidates.

### Next-generation sequencing libraries preparation

In brief, PCR products were purified by SPRI on Ampure XP paramagnetic microbeads (Beckman Coulter, USA) prior to library construction and quantified on a Qubit 2.0 instrument with the help of the dsDNA High Sensitivity Quantitation kit (Thermo Fisher Scientific). A purified PCR product (50 ng) was used for construction of libraries by means of the NEBNext® Ultra™ II DNA Library Prep Kit for Illumina (New England Biolabs, USA). KAPA UDI Primer Mixes (KAPA biosystems, cat. No. 09134336001) were employed for library indexing to minimize the risk of library cross-contamination. Illumina adapters were ligated to the resultant fragments and barcoded by PCR with primers from the NEBnext multiple oligos for the Illumina kit (New England Biolabs, cat. # E7600) and Phusion DNA polymerase (Thermo Fisher Scientific). Sequencing was performed on the Illumina MiSeq 2500 instrument (Illumina, USA). The coverage per sample was 88 to 698 thousand 150-nt paired-end reads. To map reads to the HBV genome and identify nucleotide substitutions in them, the UGENE v42.0-dev software package (Unipro, Russia) and the BWA-MEM and SAMtools tools built into it were used. To reduce the probability of mistaking sequencing errors for polymerase errors at the stage of selecting mutants, all reads of individual nucleotides (either matching the reference or different from it) with a Q-value <30 (the probability of sequencing error is >0.01%) were discarded. Further calculations were performed in Microsoft Excel 2019. All detected nucleotide substitutions that represented a deviation from the reference and were not part of the primers were designated as polymerase errors. First, for each nucleic-acid sample, the frequency of nucleotide substitutions in the final pool of amplicons was evaluated as:

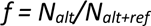

where ***f*** is the average frequency of nucleotide substitutions for all positions, ***N_alt_*** is the number of nucleotide reads that differed from the reference (total for all positions of the target), and ***N_alt+ref_*** is the total number of nucleotides read (total for all positions of the amplicon). Polymerase fidelity was then calculated from the error rate via the formula:

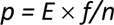

where ***p*** is the error rate, ***f*** denotes the error rate in the final amplicon population, ***E*** is PCR efficiency, and ***n*** represents the number of amplification cycles. The efficiency of each individual reaction was assessed by means of the fluorescence curve shape in the LinRegPCR software [42].

For each enzyme, the experiment was conducted in triplicate, and the resulting fidelity assessment was the averaged data.

### Parametrization of protein sequences

In our work, we employed transformer-based PLMs to parametrize mutated protein sequences into dense vectors. In particular, we utilized the encoder part of the encoder-decoder ProtT5-XL model [5], using last-layer embeddings of the encoder for parametrization. The resulting per-token sequence embeddings were next aggregated by average pooling to obtain sequence level representations. These served as input to the predictor on top of the language model embeddings.

We fine-tuned the last six layers of the ProtT5-XL encoder using the masked language modeling objective because the authors of ref. [12] argue that fine-tuning on homologs of a target protein sequence greatly improves the results of protein function prediction models, and our experiments suggested the same, though the effect was less drastic for larger models.

We chose the model implementation released in the HuggingFace library [43] and fine-tuned it with ZeRo stage 2 optimization [44] available in the DeepSpeed library by Microsoft (https://github.com/microsoft/DeepSpeed). The model was fine-tuned for 2 weeks on a server containing four NVIDIA V100 GPUs at a batch size of 1024 and a learning rate of 1e-5, with a linear decay schedule, and 200 warmup steps.

The data employed to fine-tune the model were retrieved from the UniRef100 database. Homologous sequences were extracted using jackhmmer [45] with default settings and the Klenow fragment of Taq pol (P19821) as a reference. At the time of data acquisition (January 2022), the UniRef100 database contained 280,483,851 proteins. The extracted set of homologous proteins consisted of 91,808 sequences with lengths up to 949 aa (mean length of 703.3 and a median of 875 aa).

### Regression models

In this project, we employed regression models on top of embeddings to predict functional effects of amino acid substitutions in Taq pol. For the first batch of data, we utilized the Ridge regression to capture a relation between the embeddings and the target properties. The ridge regression model was chosen for its simplicity and effectiveness in handling multicollinearity among features. We fine-tuned the α parameter by leave-one-out cross-validation, with mean absolute percentage error (MAPE) as the evaluation metric.

For the second batch of data, we switched to Gaussian Processes using the GPytorch library [46]. This decision was driven by the limitations observed in linear models, which sometimes produced extreme predictions. Gaussian Processes have the additional advantage of quantifying the uncertainty of predictions, thereby offering a more nuanced understanding of a model’s confidence in its outputs.

Each property was modeled using manually selected kernels and their respective parameters. We primarily utilized Matern or SpectralDelta kernels and adjusted the number of deltas as needed to best capture the underlying data patterns. This manual selection process ensured that the kernels were well-suited to specific characteristics of each property, thereby enhancing a model’s predictive performance.

To prepare the targets for the modeling, all values were scaled by subtraction of the mean and dividing by the standard deviation. Additionally, some targets were logarithmically transformed beforehand to stabilize variance and normalize the distribution. After predictions were made by Gaussian Processes, the density of the predictions was transformed back to the nonlogarithmic space in order to maintain interpretability. All parameters and their configurations can be found in the **Table S2**.

### Selection and validation of predictive models

Lacking direct experimental data for Taq pol, we availed ourselves of deep mutational scanning data from BlaC [47] and avGFP [48] for benchmarking, which later became a part of the ProteinGym benchmark [49]. The latter provides extensive mutational data for over 200 proteins, making it beneficial for this research field.

For model selection, we performed repeated random sampling of small training sets, which allowed us to estimate the distribution of metrics by means of the remaining data. This approach reflects our initial setting of small training sets (e.g., 18 mutants) for Taq pol. We evaluated model performance as Spearman’s correlation coefficients and a set of Top-k metrics adapted from [12].

Spearman’s correlation—a rank-based metric—was chosen to evaluate the ordinal relation between predicted and actual activity values because our primary goal was to rank mutants by functionality rather than predict exact values. Top-k metrics (k = 4, 8, 16, and 24) were used to describe the hits that exceeded the WT in terms of functionality among the top predictions. These metrics are especially useful in real-world scenarios where the selection of enhanced mutants is based on a model’s top predictions, and the extent of wet-lab experimental validation is limited.

Fine-tuned ProtT5-XL yielded results comparable to those of non–fine-tuned ProtT5-XXL, and we selected the ProtT5-XL model for protein parametrization owing to its smaller size.

### Mutational scanning at the chosen sites

The selection of the mutation set for our study was guided by several critical considerations. Firstly, we limited the number of substitutions per mutant to no more than three. This approach was adopted to avoid scenarios where a substitution can disrupt useful properties of the enzyme and this phenomenon may go undetected by our regression models. These models, designed to simulate a fitness landscape, may not accurately predict such problems owing to being trained on relatively sparse data. To refine our mutation site selection procedure, we relied heavily on insights from pertinent literature. Our literature review was instrumental in narrowing down the search space; exhaustively searching for all possible triple substitutions across all Taq pol mutants would have resulted in over 10^11^ data points. Consequently, our mutational scanning encompassed all single mutants at every position in Taq pol along with double and triple mutants specifically at aa 507, 515, 536, 540, 570, 573, 578, 586, 614, 626, 639, 670, 667, 707, 708, 728, 732, 742, 743, 747, and 783. This targeted approach enabled us to efficiently and effectively explore mutations having the potential to alter the enzyme’s properties in line with our study’s goals.

An outline of the Taq pol design and evaluation pipeline is given in **Fig. 2**.

**Figure 2.**
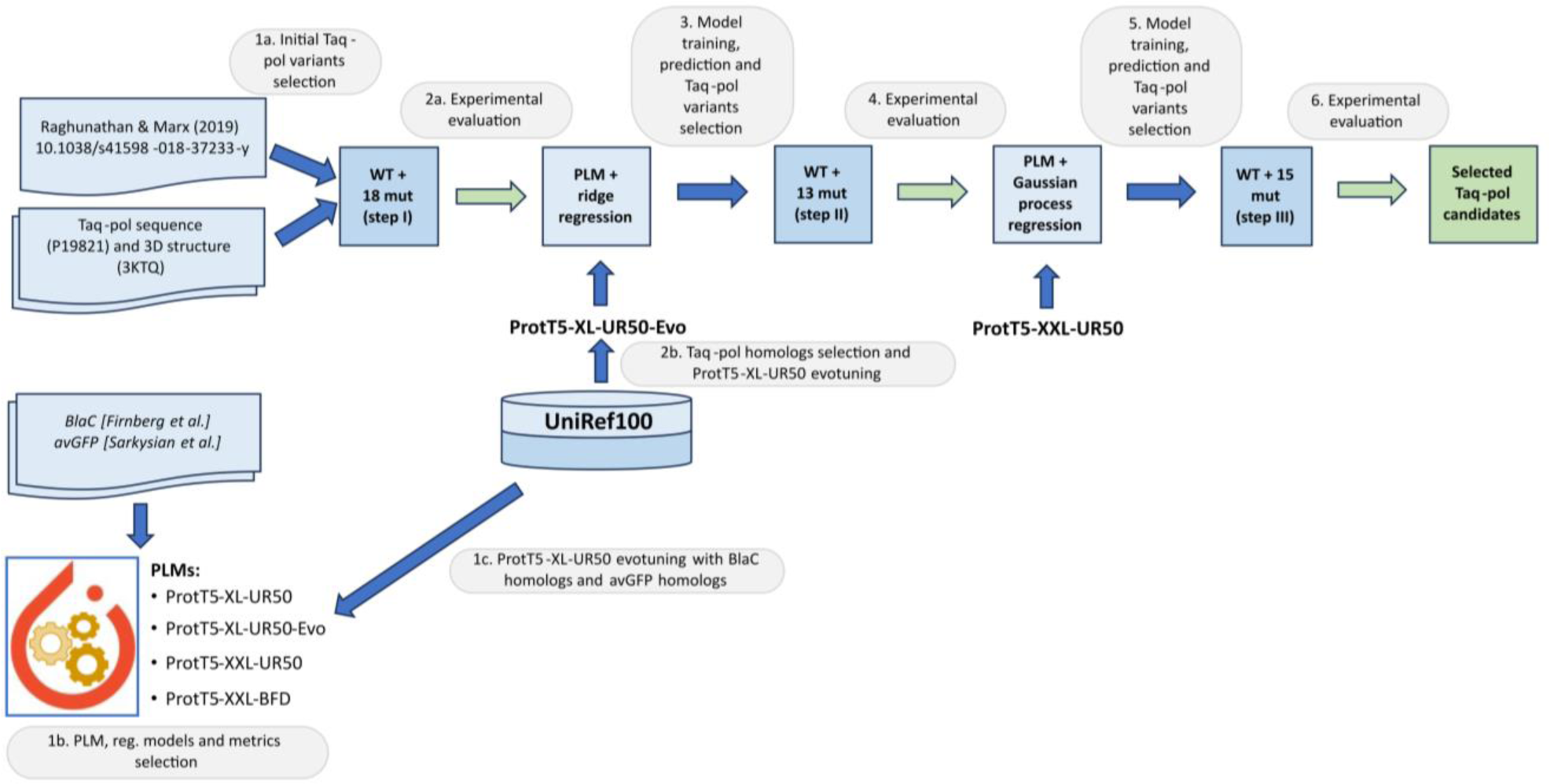
The outline of the Taq pol design and evaluation pipeline. First, the initial aa substitutions were selected for wet-lab experimental evaluation (1a), *in silico* analyses of PLMs and different regression models for protein function enhancement were performed with published mutational data (1b), and the chosen PLM was fine-tuned with Taq pol homologs (evotuned) (1c). Then, the first round of experimental evaluation was performed (2a), and the evotuned ProtT5-XL-UR50-Evo model was used to obtain embeddings of Taq pol and a vast set of its mutants with 1–3 aa substitutions (2b), after which the first regression model was built, and a new set of Taq pol mutants was selected for validation (3). After the experiments (4), the second model was obtained based on ProtT5-XXL-UR50 and Gaussian Process regression (5), and another set of Taq pol mutants was chosen for experimental assessment (6).

## Results

### Multiparametric testing of the first 18 mutant polymerases

The results of testing of the first 18 mutant enzymes in comparison with the WT enzyme are summarized in **Table 1**. For the WT, the results were comparable to those in the literature. Despite the expected loss of PCR activity in four mutants (S515F, H639A, F667A, and M747D) according to [37], in our experiments all the enzymes manifested the DNA-dependent DNA polymerase activity and the ability to perform real-time PCR with cleavage of fluorescently labeled probes. Nevertheless, in all cases, the rate of this synthesis was found to be more or less reduced as compared to the WT. PCR efficiency was quite high in all cases, 1.82 to 1.99, except for S515D (1.73) and F667A (1.59). As for mutants S515F and M747D, they were fully PCR-active.

**Table 1.**
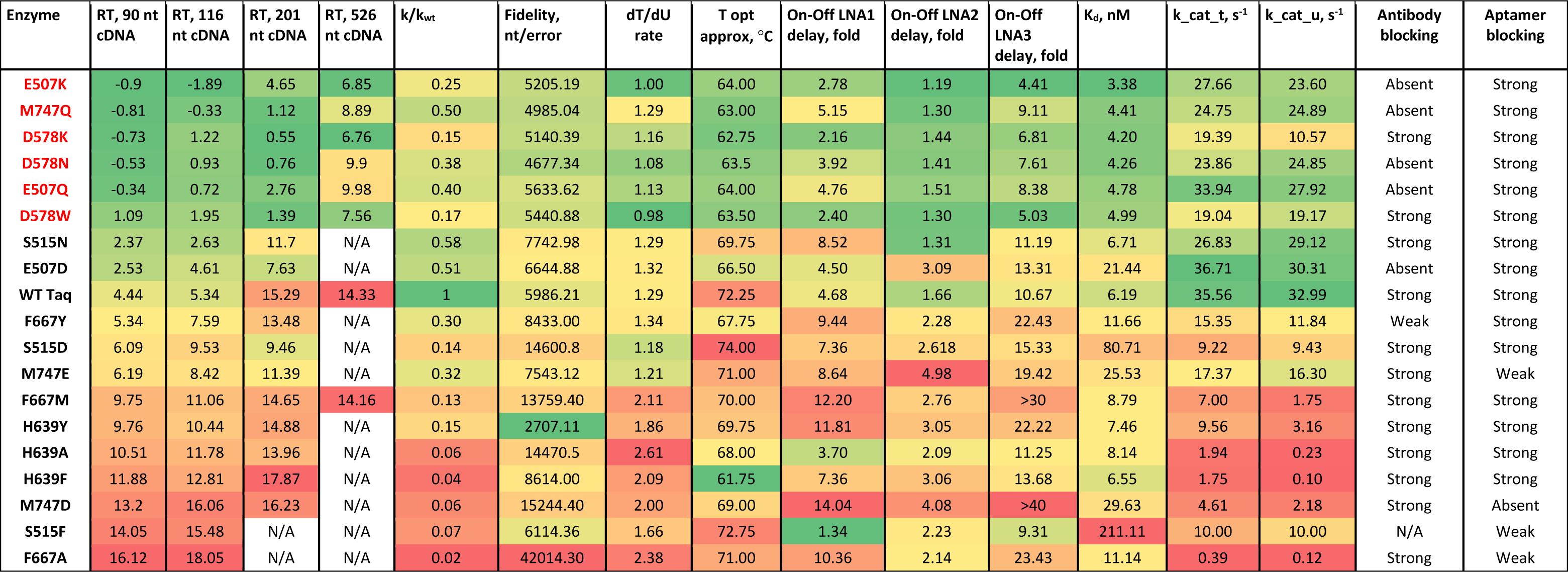
Results of multiparametric experimental testing of 18 Taq pol mutants and of the WT enzyme. Enzymes that manifested sufficient RTase activity are labeled in red. RTase activity was assessed as the difference in Cq values in the synthesis of each cDNA between a Taq pol mutant and the p66 RTase. The delay caused by the presence of LNA in a model substrate was estimated as the difference in reaction rates between a substrate containing an LNA nucleotide and the control substrate (without such a modification); nt = nucleotides

| Enzyme | RT, 90 nt cDNA | RT, 116 nt cDNA | RT, 201 nt cDNA | RT, 526 nt cDNA | k/k<sub>wt</sub> | Fidelity, nt/error | dT/dU rate | T opt approx, °C | On-Off LNA1 delay, fold | On-Off LNA2 delay, fold | On-Off LNA3 delay, fold | K<sub>d</sub>, nM | k<sub>cat</sub><sub>t</sub>, s<sup>-1</sup> | k<sub>cat</sub><sub>u</sub>, s<sup>-1</sup> | Antibody blocking | Aptamer blocking |
| --- | --- | --- | --- | --- | --- | --- | --- | --- | --- | --- | --- | --- | --- | --- | --- | --- |
| **E507K** | -0.9 | -1.89 | 4.65 | 6.85 | 0.25 | 5205.19 | 1.00 | 64.00 | 2.78 | 1.19 | 4.41 | 3.38 | 27.66 | 23.60 | Absent | Strong |
| **M747Q** | -0.81 | -0.33 | 1.12 | 8.89 | 0.50 | 4985.04 | 1.29 | 63.00 | 5.15 | 1.30 | 9.11 | 4.41 | 24.75 | 24.89 | Absent | Strong |
| **D578K** | -0.73 | 1.22 | 0.55 | 6.76 | 0.15 | 5140.39 | 1.16 | 62.75 | 2.16 | 1.44 | 6.81 | 4.20 | 19.39 | 10.57 | Strong | Strong |
| **D578N** | -0.53 | 0.93 | 0.76 | 9.9 | 0.38 | 4677.34 | 1.08 | 63.5 | 3.92 | 1.41 | 7.61 | 4.26 | 23.86 | 24.85 | Absent | Strong |
| **E507Q** | -0.34 | 0.72 | 2.76 | 9.98 | 0.40 | 5633.62 | 1.13 | 64.00 | 4.76 | 1.51 | 8.38 | 4.78 | 33.94 | 27.92 | Absent | Strong |
| **D578W** | 1.09 | 1.95 | 1.39 | 7.56 | 0.17 | 5440.88 | 0.98 | 63.50 | 2.40 | 1.30 | 5.03 | 4.99 | 19.04 | 19.17 | Strong | Strong |
| **S515N** | 2.37 | 2.63 | 11.7 | N/A | 0.58 | 7742.98 | 1.29 | 69.75 | 8.52 | 1.31 | 11.19 | 6.71 | 26.83 | 29.12 | Strong | Strong |
| **E507D** | 2.53 | 4.61 | 7.63 | N/A | 0.51 | 6644.88 | 1.32 | 66.50 | 4.50 | 3.09 | 13.31 | 21.44 | 36.71 | 30.31 | Absent | Strong |
| **WT Taq** | 4.44 | 5.34 | 15.29 | 14.33 | 1 | 5986.21 | 1.29 | 72.25 | 4.68 | 1.66 | 10.67 | 6.19 | 35.56 | 32.99 | Strong | Strong |
| **F667Y** | 5.34 | 7.59 | 13.48 | N/A | 0.30 | 8433.00 | 1.34 | 67.75 | 9.44 | 2.28 | 22.43 | 11.66 | 15.35 | 11.84 | Weak | Strong |
| **S515D** | 6.09 | 9.53 | 9.46 | N/A | 0.14 | 14600.8 | 1.18 | 74.00 | 7.36 | 2.618 | 15.33 | 80.71 | 9.22 | 9.43 | Strong | Strong |
| **M747E** | 6.19 | 8.42 | 11.39 | N/A | 0.32 | 7543.12 | 1.21 | 71.00 | 8.64 | 4.98 | 19.42 | 25.53 | 17.37 | 16.30 | Strong | Weak |
| **F667M** | 9.75 | 11.06 | 14.65 | 14.16 | 0.13 | 13759.40 | 2.11 | 70.00 | 12.20 | 2.76 | >30 | 8.79 | 7.00 | 1.75 | Strong | Strong |
| **H639Y** | 9.76 | 10.44 | 14.88 | N/A | 0.15 | 2707.11 | 1.86 | 69.75 | 11.81 | 3.05 | 22.22 | 7.46 | 9.56 | 3.16 | Strong | Strong |
| **H639A** | 10.51 | 11.78 | 13.96 | N/A | 0.06 | 14470.5 | 2.61 | 68.00 | 3.70 | 2.09 | 11.25 | 8.14 | 1.94 | 0.23 | Strong | Strong |
| **H639F** | 11.88 | 12.81 | 17.87 | N/A | 0.04 | 8614.00 | 2.09 | 61.75 | 7.36 | 3.06 | 13.68 | 6.55 | 1.75 | 0.10 | Strong | Strong |
| **M747D** | 13.2 | 16.06 | 16.23 | N/A | 0.06 | 15244.40 | 2.00 | 69.00 | 14.04 | 4.08 | >40 | 29.63 | 4.61 | 2.18 | Strong | Absent |
| **S515F** | 14.05 | 15.48 | N/A | N/A | 0.07 | 6114.36 | 1.66 | 72.75 | 1.34 | 2.23 | 9.31 | 211.11 | 10.00 | 10.00 | N/A | Weak |
| **F667A** | 16.12 | 18.05 | N/A | N/A | 0.02 | 42014.30 | 2.38 | 71.00 | 10.36 | 2.14 | 23.43 | 11.14 | 0.39 | 0.12 | Strong | Weak |

Eight enzymes showed higher RTase activity compared to the WT, and for six of the eight (two mutants in the thumb domain, three mutants in the palm domain, and one mutant in the finger domain), this improvement was substantial. The considerable increase in RTase activity in these proteins was detected both by RT-PCR and by elongation of oligo(dT) primers on a poly(rA) substrate.

The enzymes with noticeably enhanced RTase activity showed (see **Table 1**) the following:

- higher catalytic constants for both dT-containing and dU-containing dNTP mixes;
- fidelity of DNA-dependent DNA polymerase activity comparable to that of the WT or lower; a decreased (except for M747Q) dT/dU rate;
- stronger affinity (lower K_d_) for a DNA substrate as compared to the WT enzyme;
- reduced negative effects of an LNA in the model substrates, regardless of its position;
- a lowered temperature optimum of DNA-dependent DNA polymerase activity.

In all cases, the cDNA synthesis ability of mutant enzymes decreased markedly with increasing target length. dC_q_ values of all mutant enzymes versus p66 showed the highest correlation between the 90-nt and 116-nt targets (R^2^ = 0.99). With the shortest fragment, some Taq mutants outperformed the control p66 enzyme. Two enzymes failed to produce detectable 201-nt cDNA, and regarding 526-nt cDNA, it could be generated only for eight enzymes out of the 19, always with a substantial lag behind p66. The ability to process the longest target clearly correlated with catalytic constants and negatively correlated with the LNA-mediated delay and K_d_.

Some mutations led to the loss of the blocking of the enzyme by the antibody or the DNA aptamer. Namely, the M747Q mutation resulted in a loss of binding to the antibody while affinity for the aptamer was retained. In contrast, M747D led to loss of binding to the DNA aptamer but had no effect on the blocking by the antibody.

Utilizing the initial version of our model, which was trained on the dataset comprising the 19 data points, we made predictions of various properties of Taq pol mutants while specifically targeting those within a conservative trust radius of three amino acid substitutions away from the WT sequence. This constrained approach was chosen because our primary objective was to develop industrial candidate enzymes, which must meet multiple stringent requirements to function effectively. Therefore, we focused on mutations at specific sites identified in [37]. As a result, over 18 million Taq pol mutants were assessed under our prediction model.

### Selection and testing of the second and third batches of mutant polymerases

In further experiments, for resource-saving reasons, we reduced the set of experimentally estimated parameters: 1) excluded the LNA-2 assay, which simulated the incorporation of an LNA nucleotide into a newly synthesized strand (this was outside the scope of our purposes); 2) excluded the LNA-1 assay as redundant in relation to the LNA-3 assay (the same characteristic and a less pronounced effect); excluded K_cat_ (dT) and K_cat_ (dU) measurement because of low throughput (the same features of Taq pol mutants were indirectly evaluated by higher-throughput assays); 4) skipped the assessment of the temperature optimum, which in our experiments showed too high variation. Of the remaining parameters, RTase efficiency on 90-nt and 116-nt templates, the effective reaction rate constant, fidelity, PCR efficiency, the dT/dU rate, K_cat_ (dT), and K_cat_ (dU) (based on the experimental results obtained on the first batch of enzyme variants), and probability of enzyme blocking by the antibody or by the DNA aptamer were chosen as independent parameters for model training. The LNA-3 substrate delay, RTase efficiency on the 201-nt and 526-nt templates, and RTase activity measurements with the oligo(dT)-poly(rA) complex were not used for the training (because these characteristics could not be reliably evaluated by our methods in all enzymes) but were monitored in the wet-lab experiments.

To assess predictive power of the newly developed algorithm and for its possible further refinement, we selected and synthesized another 13 mutant Taq pols on the basis of both the model predictions and literature data. The logic of their choice was as follows:

- to include new sites for a single substitution as compared to those used in the initial model training; the new substitutions had to have a positive impact on the quality of subsequent predictions (R660S and D732N);
- to include two- or three-substitution mutants expected to evolve in opposite directions according to the model predictions: enhanced RTase activity (E507K-D732N, D578H-M747Q, E507Q-S515N-D578S, E507Q-D578S-I614M, E507Q-D578S-R728Q, D578S-R728Q-M747Q, and D578S-R728Q-M747V) and higher fidelity (V586N-M747T, S515N-M747S, S515D-K540M, and F667I-R728N-M747S);
- to include some mutants for which there were literature data on the properties of interest to us (R660S: enhanced allele-specificity and Sanger sequencing quality [50] [51] and D732N: stronger RTase (also predicted by our model) and strand displacement activities [24]).

According to the obtained experimental findings, the predictive model was refined and a third batch of mutant enzymes was prepared for its adjustment; these enzymes included additional new substitution positions and combinations thereof. This batch contained 15 enzymes. For eight of them (E507K-D578E, R573K-D578N, D578N-V783I, D578Q-D732N, E708Q-D732N, E507R-R573K-E708Q, E507K-A570G-M747Q, and E507K-A570T-M747Q) RTase activity enhancement was predicted by the model. For the other seven, literature data about various useful properties were available, e.g., increased allele specificity (R660V [27] and E507K-R536K-R660V [28]), cold sensitivity (I707L and E708D [31]), resistance to PCR inhibitors (E708L [21] and E742K-A743R [22]), and resistance to PCR inhibitors along with concurrent cold sensitivity (E626K [31]).

When selecting candidates for enhancing RTase activity, we were guided by the following criteria: K_cat_ (dT) and K_cat_ (dU) at least 20 nM, PCR efficiency not less than 1.85, the dT/dU rate not more than 1.5, the error rate not greater than 1/3000, the effective reaction rate constant ≥0.2 of the WT, and the probability of no blocking with the antibody or DNA aptamer not more than 0.1. The remaining characteristics were disregarded in the selection. Results of the testing of the second and third batches of enzymes are given in **Table 2**.

**Table 2.**
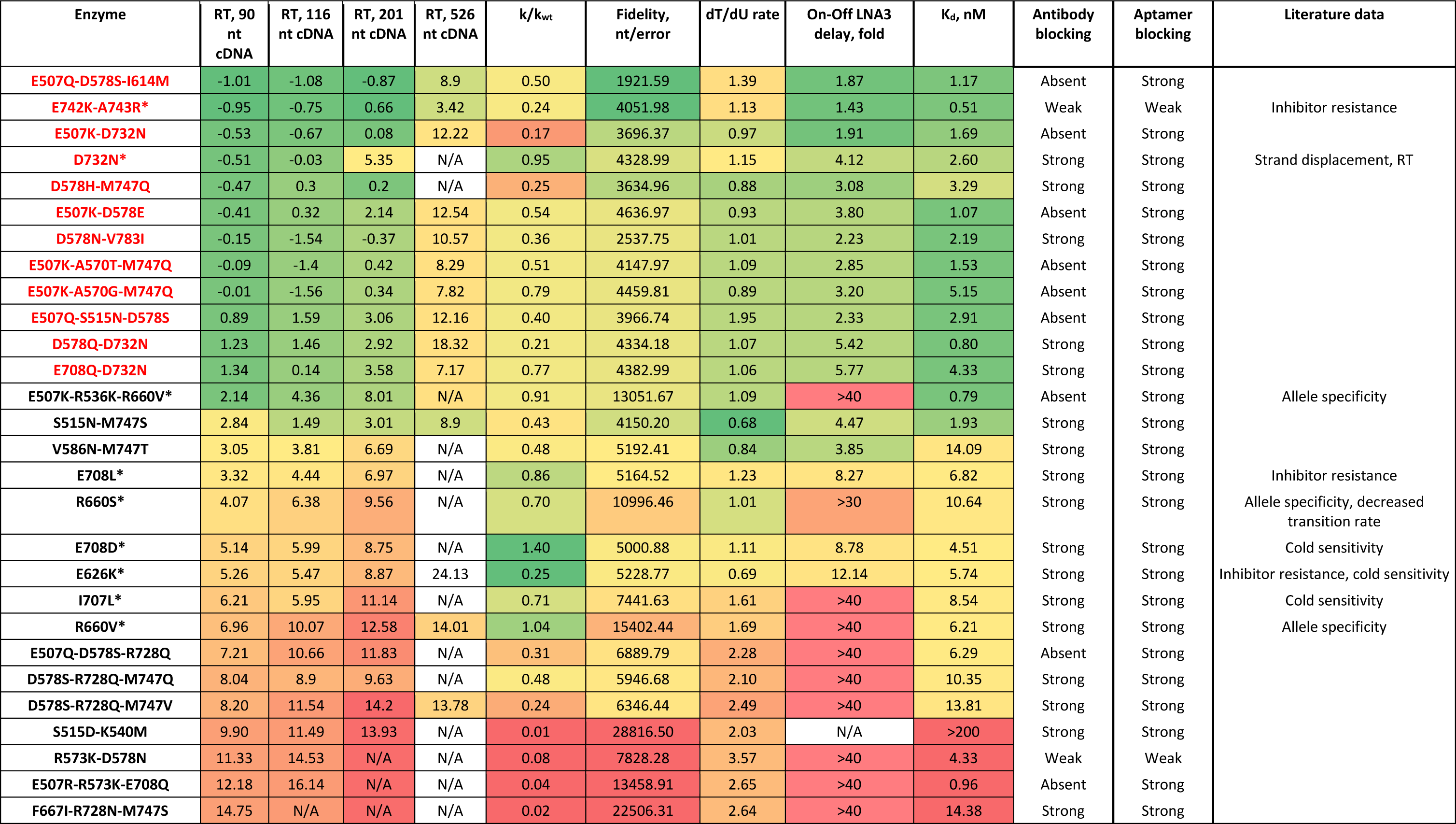
Results of multiparametric experimental testing of 28 Taq pol mutants. Enzymes that exerted sufficient RTase action are labeled in red. Enzymes selected based on literature data are marked with asterisks. RTase activity was assessed as the difference in Cq values in the synthesis of each cDNA between a Taq pol mutant and the p66 RTase. The delay caused by the presence of an LNA nucleotide in a model substrate was estimated as the difference in reaction rates between the substrate containing LNA and the control substrate (without such a modification).

In **Fig. 3**, Spearman’s correlations between the parameters experimentally measured for all 46 Taq pol mutants and the WT are presented.

**Figure 3.**
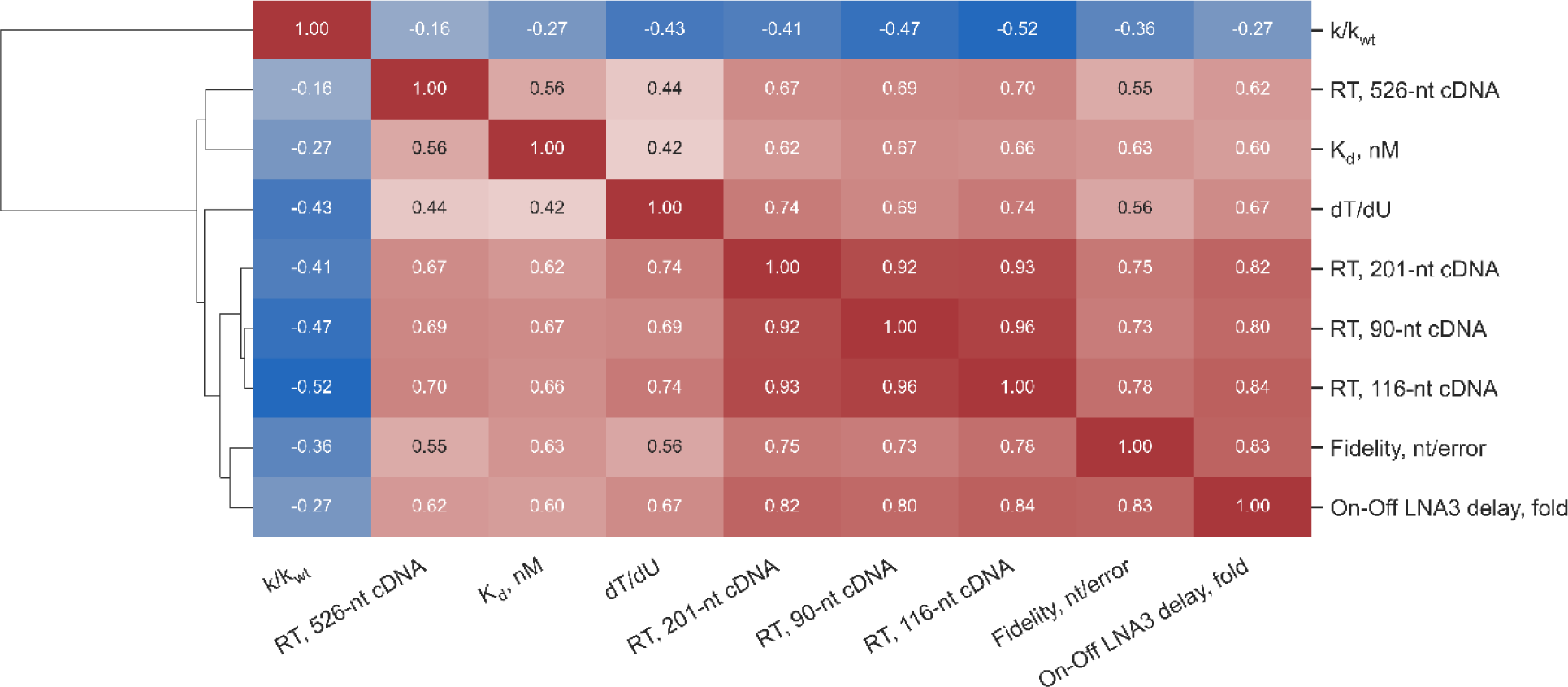
Spearman’s correlations between the parameters measured for 46 Taq pol mutants and the WT enzyme. The color scale varies from deep blue for highly negative correlation coefficients to red for highly positive ones.

As one can see, the correlations between the characteristics of mutant enzymes identified in the first batch persisted in subsequent ones: the enhancement of RTase activity was accompanied by a general trend toward a diminished dT/dU rate, slightly lower fidelity, stronger K_d_ values, and reduced negative effects of an LNA nucleotide (with the exception of D732N), whereas the enhancement of fidelity followed the opposite trend.

From **Fig. 4**, it can be deduced that in this trend, any of the enzyme groups does not differ from the total collection of tested enzymes. Some enzymes showed improved RTase activity on longer templates as compared to the best enzymes from the first batch.

**Figure 4.**
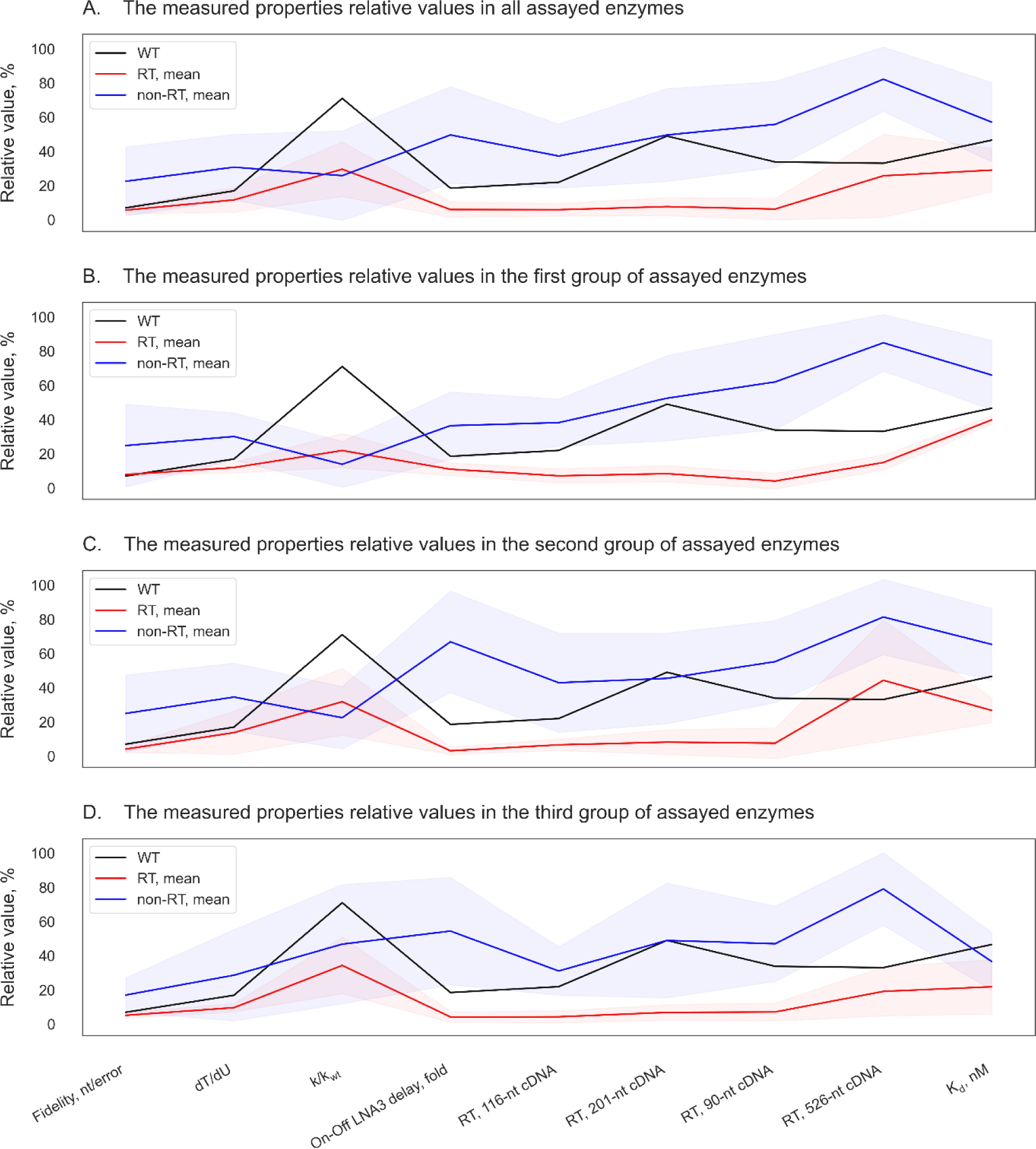
Parallel coordinate plots of the assayed Taq pol mutants and of the WT enzyme. Mean relative values of properties of non-RT enzymes (lacking appreciable RTase activity) are highlighted in blue, and the mean relative values of properties of RT enzymes (having substantial RTase activity) are shown in red. The shaded area denotes standard deviation ranges. Relative values of the WT enzyme’s properties are presented as the black lines.

Overall, the correlation between predicted and experimentally confirmed RTase activities for all enzymes was quite high (see **Fig. S2**). Unexpectedly, both mutants containing substitutions in codon 728 did not exert sufficient RTase action. Vice versa, mutant E742K-A743R, predicted to have low RTase activity, manifested its enhancement.

Again, the predictive model was refined based on the addition of the new experimental findings.

Altogether, our search identified 18 Taq pol mutants with substantially enhanced RTase activity as compared to the WT. In all cases, the cDNA synthesis ability of mutant enzymes declined markedly with increasing target length, and this feature disadvantageously distinguished them from native RTases. We hypothesized that the sharp drop of RT efficacy may be due to the extremely low strand displacement activity distinguishing Taq pol mutants from mesophilic RTases. Nonetheless, this hypothesis was refuted when the RTase capacity of two enzymes from our collection was compared: D732N, which has a pronounced strand displacement activity, and E507Q-D578S-I614M, which does not possess such an activity. When synthesizing longer cDNAs, E507Q-D578S-I614M reproducibly outperformed D732N (data not shown). Another reason could be that we carried out RT at nonoptimal temperature for this activity of a given enzyme (we did not test this explanation because this would require major redesign of our assays). In this regard, when analyzing the second and third batches of enzymes, we continued to separately monitor the performance of mutants on four RNA templates of different lengths as four independent characteristics and to utilize two of them for training the model.

### Fidelity of DNA-dependent DNA-polymerase activity

Values of overall fidelity of mutant enzymes and of the WT are listed in **Tables 1** and **2**, and a more detailed view (absolute and relative frequencies of different transitions and transversions) is given in **Fig. 5**, **Table S3**, and **Fig. S3**.

**Figure 5.**
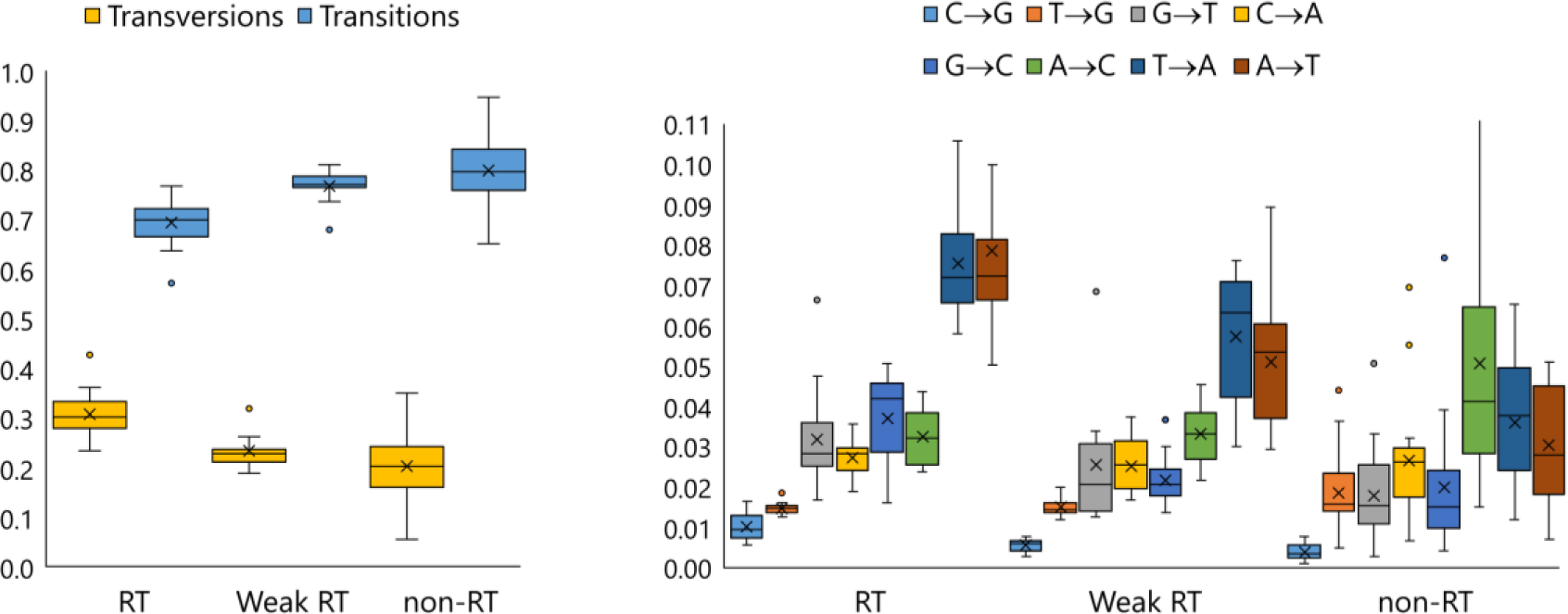
Box-whisker plots representing relative summarized frequencies of transitions and transversions (left) and relative frequencies of specific transversions (right) in DNA-dependent DNA polymerase–driven synthesis by enzymes with sufficiently enhanced RTase activity (RT, n = 18), enzymes with extremely low RTase activity (non-RT, n = 19), and enzymes intermediate in terms of this characteristic, including the WT (Weak RT, n = 11).

This parameter varied widely in the total collection of enzymes. For all enzymes with enhanced RTase activity, it a) was slightly reduced compared to the WT but exceeded the values known for HIV-1 RTase [52], and b) featured an increase in ratios of transversions to transitions, especially substitutions A->T and T->A. The increase in accuracy followed the opposite trend: it matched an increase in the proportion of transitions (R^2^ = 0.73). Not only the overall frequency of transitions and transversions (see **Table S3**) but also the proportion of various substitution types varied among the enzymes, in some cases following rather different patterns (see **Fig. S3**).

### Taq pol mutants with enhanced RTase activity in single-tube TaqMan RT-PCR

The 12 selected candidate enzymes were tested as described in the Methods section (**Fig. 6**).

**Figure 6.**
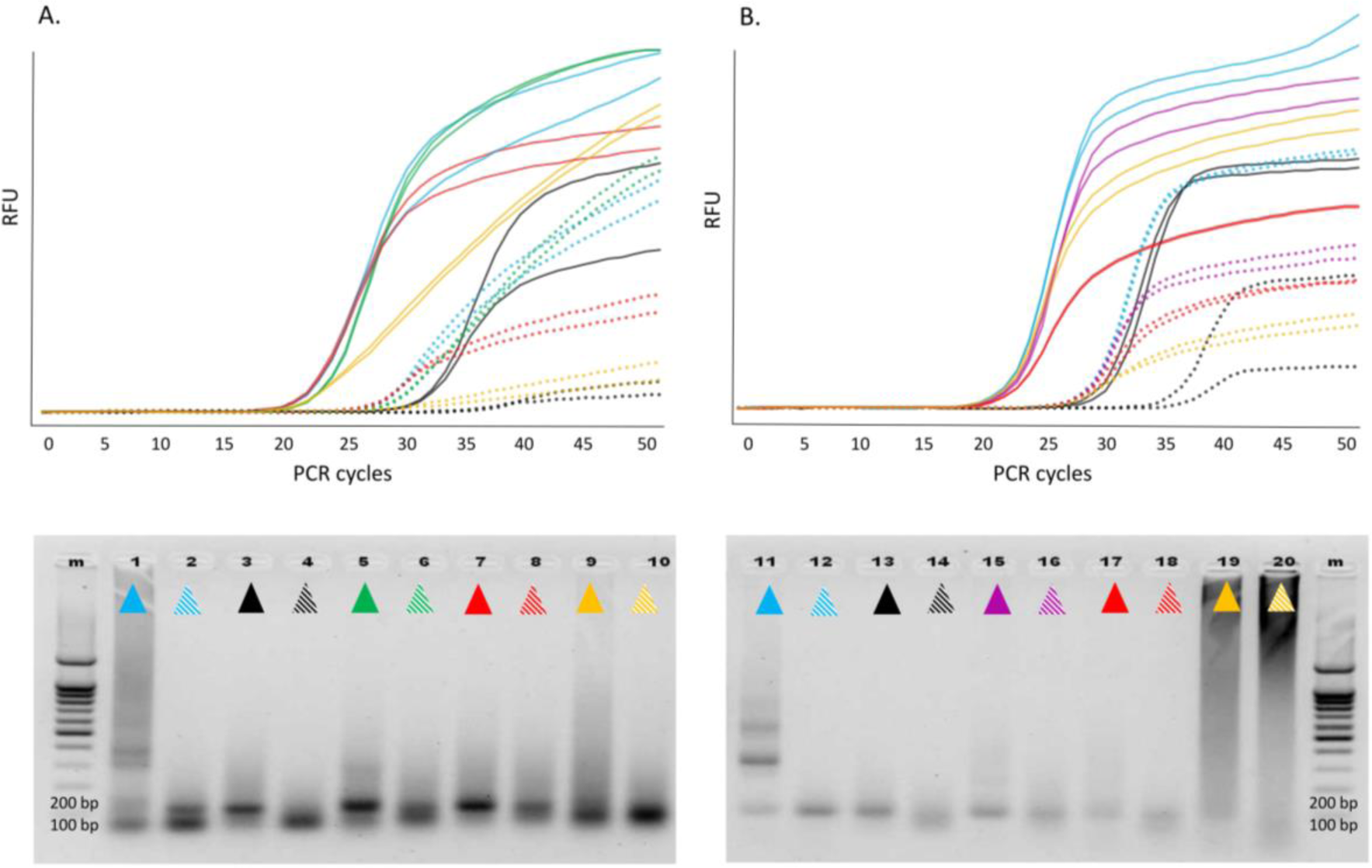
Above: Real-time RT-PCR involving various Taq pol mutants in single-enzyme reactions with a synthetic transcript containing the 116-nt hMPV sequence (A) and the 90-nt HPIV sequence (B). Color indication: WT Taq pol, WT/p66 mix, E507Q, D578N, E507Q-D578S-I614M, E507K-A570G-M747Q. For each transcript, two dilutions were analyzed, of which the second (dashed lines) is 1/100 of the first (solid lines). Below: 2% agarose gel electrophoresis of the reaction products (one well per dilution, the color indication as above). Left, hMPV; right, HPIV. M: marker. RFU, relative fluorescence units.

The findings were interpreted with respect to two parameters: C_t_ meant integral efficiency of RT and cDNA amplification, and the amplitude and shape of the fluorescence curve reflected the 5′ exonuclease activity resulting in the cleavage of the labeled probe. All the 12 candidates expectedly outperformed the WT as RTases. As compared to the p66/WT mixture, all these candidates yielded comparable or even lower C_t_ values, indicating that sensitivity was comparable between the single-enzyme mode and two-enzyme mode of target RNA detection. The shapes and amplitudes of the fluorescence curves as well as PCR efficiency of the generated cDNA in the single-enzyme reactions were comparable to those of the control two-enzyme reactions. Nonetheless, in some single-enzyme reactions, we observed a high level of low-molecular-weight and/or high-molecular-weight off-products, which negatively affected the shape and slope of the kinetic curve, especially during HPIV2 RNA detection. In our experience, this phenomenon is characteristic of enzymes with strong K_d_ and can be improved by optimizing the reaction conditions. Indeed, an increase in KCl concentration and pH of the reaction mixture led to a reduction in the amount of off-products, an improvement in the shape of the kinetic curves, and clear-cut electrophoretic bands when enzymes E507K-D732N and E507Q-D578S-I614M were used (data not shown).

## Discussion

The fact that WT Taq pol has *in vitro* RTase activity has been known for a long time [53] [54]. In numerous papers, mainly in the last decade, it has been demonstrated that this activity can be significantly enhanced by modification of reaction conditions or by introduction of amino acid substitutions at various positions (see refs. above). In our study, we deliberately limited the number of analyzed amino acid positions, thereby radically narrowing the space of possible mutants. Nevertheless, our search—involving a predictive model based on data from multiparametric enzyme testing—identified Taq pol mutants having considerably boosted RTase activity combined with a set of other desirable characteristics. In our experiments, this enhancement strongly correlated with broadened substrate specificity, including a greater ability to utilize “unnatural” substrates, in agreement with other articles in the field.

Judging by our results, the dissociation constant of an enzyme toward a DNA substrate is a key parameter responsible for the mechanism underlying the expansion of substrate specificity: it was mutants with stronger K_d_ that showed such expansion. Moreover, although K_d_ was not our criterion for the selection of candidates for wet-lab experiments, the selection for K_d_ actually occurred coupled with the selection for RTase activity. A simple explanation is that the stronger binding of mutant enzymes to various substrates enables reactions with these substrates, in contrast to WT Taq pol, for which such reactions are inhibited/prevented by the weak affinity for the substrates. On the other hand, our analysis of the same set of experimental data suggests that “1D” selection aimed at enhancing K_d_ may not be effective enough to find the best candidates for different biotechnological tasks. This is because effects of different “K_d_-strengthening” mutations on features of interaction with one or another type of substrate were different even among enzymes with high DNA-dependent DNA polymerase activity. For example, E708D has stronger K_d_ compared to the WT but showed no improvement of RTase activity, whereas V586N-M747T has weaker K_d_ but manifested moderately increased RTase activity and better tolerance to an LNA substrate. This is not surprising because the effects of these mutations on enzyme–substrate interactions can have dissimilar mechanisms. For instance, the thumb domain, which includes E507, is involved in interactions with the upstream duplex of an overlapping substrate [55]. E507 is situated in the primer–template-binding site where mutations are expected to modulate DNA-binding affinity. E507K stabilizes the Taq pol–DNA binary complex by forming additional contacts with the distal portion of the primed template [56]. S515 is important for α-helix stabilization and makes the nucleic-acid–binding motif more robust [36]. D578 (palm) comes into contact with a template strand [37]. E742K and M747K (finger) could be structurally responsible for the formation of a salt bridge with negatively charged template phosphodiester groups located close to aa 739 and 747 or with a nucleotide’s triphosphate group close to aa 817 [34]. M747K introduces an additional positive charge near the negatively charged RNA template backbone thereby possibly helping to accept an unnatural substrate by enhancing binding. In turn, D732N (finger) seems to be at a distance from primer and template strands in crystal structures [24], and its participation in enzyme–substrate interactions is not fully understood. I614 (finger) contacts an incoming nucleoside-5′-O-triphosphate. Ref. [57] indicates that Taq pol tolerates amino acid substitutions at position I614 and that such mutant enzymes retain activity similar to that of the WT enzyme, but fidelity is often low. In their experiment, however, nonhydrophilic substitutions, including I614M, did not alter the error rate during DNA synthesis. The observed influence of our tested amino acid substitutions on Taq pol RTase activity can be illustrated with a biplot of a partial least squares (PLS) model (**Fig. S4**). The PLS model was trained to predict on the basis of mutation data whether enzymes have RTase activity. When we analyzed some mutations’ frequencies in Taq pol mutants predicted to possess the RTase activity, we noticed that the majority of these proteins contain at least one of such mutations as E507K or E507R, E742Q or E742M or E742H, M747K, I707R or I707K, A570K or A570R (see **Table S4**).

Of note, the accuracy of the predictive model may be affected by assays of certain enzyme characteristics in wet-lab experiments. We focused on methods for assessing the characteristics that could be experimentally evaluated over a wide range of values, ranging from negligible to high, as opposed to categorical variables (all-or-none). Accordingly, the methodology — that we applied to the estimation of RT activity via a comparison of RT efficiency among specific RNA templates of different lengths and structures by real-time RT-PCR—turned out to be more informative than traditional direct recording of reaction kinetics through elongation of oligo(dT) primers on a poly(rA) substrate. The latter option helped us to distinguish the WT from the enzymes with sufficiently enhanced RT activity but not to reproducibly stratify them among themselves (data not shown). This could be because the optimum temperature for RTase activity of the tested Taq pol mutants exceeded 60 °C: the temperature at which we failed to obtain good-quality kinetic curves with oligo(dT) primers and the poly(rA) substrate because of the low melting temperature of their duplexes. We should emphasize that the RT and PCR conditions we used had default settings, i.e., we did not adapt them to the tested enzymes; this approach could improve the quality of the obtained data. In our experiments, however, WT Taq pol was able to synthesize cDNA longer than 500 nucleotides (albeit with low efficiency), whereas, for example, in refs. [35] [37], it failed to extend a bound DNA primer strand beyond 2–7 nucleotides on some RNA template.

Our attempts to leverage recent advancements in protein modeling involving PLMs were thwarted by hardware-related constraints. Although we managed to fine-tune only the last six layers of ProtT5-XL, we anticipate that the method’s efficiency may be improved through full-model fine-tuning or the use of larger PLMs. Parameter-efficient fine-tuning aroused considerable interest in recent years owing to its potential to facilitate fine-tuning of larger models, such as ProtT5-XXL, on the same hardware [58]. Authors of ref. [59] provide evidence that low-rank adaptation (LoRA) procedures enhance the ability of PLMs to navigate mutational fitness landscapes, indicating that parameter-efficient fine-tuning may further improve model performance without extraordinary computational resources.

The use of more advanced or specialized architectures may also enhance our predictive capabilities. Multimodal models like ProstT5, which take both sequence and 3D structure data as inputs, can provide more informed predictions. Although 3D structural information is implicitly present in sequence-derived embeddings, explicit incorporation of structural data can provide additional insights. Moreover, ProteinNPT, as described in ref. [49], learns joint representations of full input batches of protein sequences and associated property labels, thereby making possible a prediction of single or multiple protein properties, novel sequence generation via conditional sampling, and iterative protein redesign cycles through Bayesian optimization.

When proteins are designed, it is crucial to consider multiple properties, especially in industrial settings. As a mutation radius increases, it quickly becomes impossible to exhaustively explore the search space thus necessitating a search for algorithms that prioritize candidate mutations for inference. Discrete optimization methods such as genetic algorithms or Markov Chain Monte Carlo, as described in [12], are required for navigating the vast search space and exploring enhanced enzymes that are a couple more mutations away from the WT. This multicriterion optimization is computationally challenging, particularly because some properties entail tradeoffs. Finding an optimal balance under such circumstances is an unsolved problem that prevents achieving desired characteristics in industrial applications. Furthermore, restricting the number of amino acid substitutions analyzed per protein to three is of course a serious limitation here, which definitely lowers the probability of finding an optimal enzyme. A number of examples are known where the best properties of a bioengineered Taq pol modified to enhance RTase function have been achieved via introduction of more amino acid substitutions, e.g., TaqM1 (L322ML349M-S515R-I638F-S739G-E773G) [35] or RT-KTq (L459M-S515R-I638F-V669L-M747K) [60]. Consequently, we plan to address this problem in our future research.

Another important topic is zero-shot methods for selecting initial datasets when experimental data are unavailable. These procedures typically are based on the likelihood of a sequence according to a PLM. Authors of ref. [61] provide details on how such approaches can be implemented effectively to identify promising candidates for initial testing, thereby reducing the experimental burden and accelerating discovery. Although this strategy is outside the scope of our study (because we relied on literature data), it is crucial for similar projects in general.

Understanding the “linkage” of effects of individual mutations or combinations thereof on several characteristics of mutant polymerases allows for predictions of the properties of enzyme variants for which these characteristics are unknown or unpublished. For example, the I704L mutation described in the literature as leading to cold sensitivity, in our experiments (in full accordance with the predictive model) also led to weaker K_d_, an RTase activity lower even compared to the WT, diminished efficiency of dUTP incorporation, and low tolerance to a hairpin LNA-containing substrate. All three mutants possessing increased allele specificity according to literature data (R660S, R660V, and E507K-R536K-R660V) expectedly showed elevated fidelity and a greater delay in synthesis on the LNA templates as compared to the WT enzyme. At the same time, a downside of this “linkage” may be unexpected “side effects” of selection for some useful property. In our work, these were pronounced attenuation of the negative influence of an upstream LNA nucleotide and a decrease in a temperature optimum during selection for greater RTase activity. We have no doubt that the effects of amino acid substitutions and their combinations on other useful properties of enzymes, for example, tolerance to PCR inhibitors, DNA lesion bypass, or the ability to incorporate fluorescently labeled monomers, can be connected and predicted *in silico*. This accomplishment should make it possible to rationally design an enzyme with a preselected combination of properties without the need to validate *in vitro* a huge number of candidates.

It is noteworthy that in our assays, all the tested enzymes were able to cleave 5′-fluorescently labeled probes, albeit with different signal amplitudes. Although this was a desirable outcome and a criterion for candidates to be selected, the absence of mutations dramatically affecting 5′-nuclease activity prevented us from using this parameter to train the predictive model. Therefore, we cannot rule out that some of the candidates selected with it would be devoid of this activity.

Important limitations of the study should be mentioned. Firstly, some key characteristics (e.g., the Michaelis constant, processivity, and K_d_ toward dNTPs) were not evaluated at all. Secondly, some identified patterns could be attributed to specific reaction conditions. Thirdly, we evaluated only the fidelity of DNA-dependent DNA polymerase activity because we could assess this characteristic in the entire set of mutant enzymes. Regarding the fidelity of RNA-dependent DNA polymerase–driven synthesis, it was determined only in a small number of enzymes (data not shown). Fourthly, we established a purely empirical relation of individual mutations or their combinations with properties of the enzyme without examining the physicochemical mechanisms underlying the effects of these mutations. For example, the target length–dependent decline of cDNA synthesis efficiency in all the analyzed Taq-derived RTases may be associated with higher processivity of the latter, strand displacement activity, a lesser degree of product inhibition, or other factors or combinations thereof. The observed decreased synthesis rate of the polymerase on hairpin LNA-containing templates may be subject to different interpretations too (see [62] [63] [64]).

### Conclusion

Through a screening of a collection of 47 mutant Taq DNA polymerases—29 of which were selected via our proposed strategy of multiparametric rational design—we were able to identify 18 enzymes that possess substantially enhanced RTase activity as compared to the WT enzyme; 12 of these Taq pol mutants were selected by our AI-based algorithm. The analyzed mutants contain amino acid substitutions affecting 20 positions in all three structural domains of Taq pol. As predicted by our algorithm and subsequently confirmed experimentally, the RTase activity enhancement tends to be accompanied by stronger K_d_ values, moderately decreased fidelity, and greater tolerance to noncanonical substrates such as nucleic acids containing dUTP and/or LNA nucleotides. Some Taq candidates were effective in single-enzyme RT-PCRs involving cleavage of fluorescently labeled probes or an antibody- or aptamer-mediated hot start. Therefore, they can provide the basis for the creation of new tools for real-time RT-PCR technologies such as pathogen RNA detection or gene expression analysis. We regard our results as proof-of-concept data, not as a final solution even in relation to the problem in question, and we do not rule out the possibility of optimizing our approach toward an analysis of combinations of more mutations. Besides, when new enzymes are bioengineered to solve specific biotechnological problems, it must be borne in mind that enhancing one function may entail inevitable and multidirectional alterations of other functions, sometimes in an unpredictable manner. Nonetheless, deep learning models proved to be valuable for guiding our selection of mutations, thereby highlighting good potential of AI-driven approaches in enzyme engineering especially in settings with a relatively small number of experimental studies.

## Contributions

Conceptualization and Methodology: NER, DVA, DNS, YET, MKI, and GBP; Validation: NER, IMY, YET, and MKI; Software: NER, IMY, and EVS; Formal analysis: NER, IMY, EVS, DVA, VNT, and MKI; Investigation: MKI, YET, EVB, LOB, SOB, OST, NSG, DVP, MOA, and AAA; Visualization: MKI, YET and DVA; Writing - Original Draft: MKI, NER, DVA, IMY, and YET; Writing - Review & Editing: MKI, NER, DVA, IMY, and YET; Supervision and Project administration: MKI and DNS; Funding acquisition: MKI and DNS.

## Conflicts of interest

All authors declare that they have no conflicts of interest.

## Acknowledgements

This work/article was partially supported by the Research Program at the MSU Institute for Artificial Intelligence. The reagents and experimental studies were supported by AO Vector-Best.

## Notes

https://huggingface.co/datasets/nerusskikh/taqpol_insilico_dms

### Competing Interest Statement

The authors have declared no competing interest.

### Summary of Updates

The link to online resource was added: https://huggingface.co/datasets/nerusskikh/taqpol_insilico_dms

