## Supplementary material for "Enhancing the Reverse Transcriptase Function in Taq Polymerase via AI-driven Multiparametric Rational Design": Table S1

**Table S1. Oligonucleotides used in the study.**

**1)	Hairpin templates. Bold denotes self-complementary regions. (+T) denotes LNA-T modification. SIMA, Dichloro-diphenyl-fluorescein.**

H-Kd (for Kd determination)

5’-GCTATGCGTGAGCGCT(T-SIMA)TTGCGCTCACGCATAGC

20_60 (for kCatT and kCatU determination)

5’-AAACTCAAGCCTTCCACATATTCTATCCACGACTCTTTACAGGTAGTATAATCTCACTATCGGACTGACTGCGTGAGCGCTTTTGCGCTCACGCAGTCAGTCCG

OnOff1_7G-C_long1 (for kinetic analyses)

5'-ATCTCGCTGTACTCACCGGTTCCGCAGATCGACATACTTATTAACTTATCTATCCTCTTACTTCATCTCATCGATACTTTTGTATCGATGAGATGAAG

LNA 0-3 (for the analysis of LNA-associated effects)

LNA-0 5'-CACTCCACAGGTCGCATTGCTAGCTGGTTAGAGACTAGTAGATCGGAAGAGCCATGAACTCCUACACTCTACGGCTCTTCCGATC

LNA-1	5'-CACTCCACAGGTCGCATTGCTAGCTGGTTAGAGACTAGTAGATCGGAAGAGCCATGAACTCCUACACTCTACGGCTCTTC(+C)GATC

LNA-2	5'-CACTCCACAGGTCGCATTGCTAGCTGGTTAGAGA(+C)TAGTAGATCGGAAGAGCCATGAACTCCUACACTCTACGGCTCTTCCGATC

LNA-3	5'-CACTCCACAGGTCGCATTGCTAGCTGGTTAGAGACTAGTAGATCGGAAGAGCCATGAACTCCUACACTCTACGGCTCTTCCGA(+T)C

**2)	Primers and fluorescently labeled probes for RT and PCR. (+T) and (+C) denote LNA-T and LNA-C modifications. ROX, 6-carboxyl-X-Rhodamine. BHQ2, Black Hole Quencher-2. R, purine (A or G). F, forward primer. R, reverse primer. P-probe.**

HBV-F	5'-TGTCTGCGGCGTTTTATCA

HBV-R	5'-AGGACAAACGGGCAACATAC

HBV-P	5'-ROX -CATCCTGCT(BHQ2)GCTATGCCTCATCTTCTTRTTGG-p

HPIV2-F	5'-AAAGAGCAAGAGGCAACCATA

HPIV2-R	5'-CACACCTGGGATTACACCA

HPIV2-P	5'-(ROX)-TCT(+C)CGACCCA(T-BHQ-2)GCAATACTCCATTCTCTC-p

hMpV-F	5'-CTCTACAGGCAGCAAAGCA

hMpV-R	5'-CATTATATTGTTAGATGACCTGGCAA

hMpV-P	5'-(ROX) CC(+C)(+C)ACCT(T-BHQ-2)AG(+C)AGTGTTTGA(+C)C-p

RhV-F	5'-CTAGCCTG(+C)G(+T)GG

RhV-R	5'-GAAACACGGACACCCAAAGTA

RhV-P	5'-(ROX)TC(+C)GGCC(+C)C(T-BHQ-2)GAATGTGGCTAA-p

HIV1-F	5’-GACTCTGGTAACTAGAGATCCCTCAGA

HIV1-R	5'-CTTGGTGTCTCTTATGTCTATCTTTTG

HIV1-P	5’-(ROX)-TCTCTAGCAGTGGCGCCCGAACA-(BHQ2)
