## Supplementary material for "Enhancing the Reverse Transcriptase Function in Taq Polymerase via AI-driven Multiparametric Rational Design": Table S3

| **Enzyme** | **Fidelity** | **C>G** | **T>G** | **G>T** | **C>A** | **G>C** | **A>C** | **T>A** | **A>T** | **C>T** | **G>A** | **A>G** | **T>C** | **TV** | **TS** | **TV/TS** | **%TS** | **%TV** |
| --- | --- | --- | --- | --- | --- | --- | --- | --- | --- | --- | --- | --- | --- | --- | --- | --- | --- | --- |
| **E507Q-D578S-I614M** | 1921,59 | 1,28 | 1,53 | 4,57 | 3,27 | 4,47 | 2,67 | 10,58 | 14,46 | 11,10 | 13,89 | 15,58 | 16,61 | 42,83 | 57,17 | 0,75 | 0,0298 | 0,0223 |
| **D578N-V783I** | 2537,74 | 1,62 | 1,83 | 6,64 | 3,06 | 2,96 | 2,46 | 9,68 | 7,92 | 5,10 | 13,24 | 18,00 | 27,47 | 36,18 | 63,82 | 0,57 | 0,0252 | 0,0143 |
| **H639Y** | 2707,11 | 0,09 | 0,47 | 0,25 | 0,66 | 0,41 | 1,47 | 1,37 | 0,67 | 7,56 | 17,52 | 25,86 | 43,68 | 5,39 | 94,61 | 0,06 | 0,0350 | 0,0020 |
| **D578H-M747Q** | 3634,96 | 0,58 | 1,31 | 2,83 | 2,80 | 5,06 | 4,26 | 6,17 | 11,44 | 12,00 | 13,38 | 16,49 | 23,67 | 34,45 | 65,55 | 0,53 | 0,0180 | 0,0095 |
| **E507K-D732N** | 3696,37 | 0,81 | 1,45 | 3,21 | 2,90 | 4,61 | 3,72 | 6,68 | 9,98 | 10,73 | 13,89 | 16,94 | 25,06 | 33,37 | 66,63 | 0,50 | 0,0180 | 0,0090 |
| **E507Q-S515N-D578S** | 3966,74 | 0,54 | 1,37 | 2,91 | 3,17 | 4,88 | 3,81 | 5,86 | 8,78 | 10,36 | 14,19 | 17,35 | 26,78 | 31,32 | 68,68 | 0,46 | 0,0173 | 0,0079 |
| **E742K-A743R** | 4051,98 | 1,56 | 1,44 | 4,75 | 3,55 | 2,95 | 2,37 | 9,95 | 7,12 | 6,45 | 18,87 | 14,93 | 26,06 | 33,69 | 66,31 | 0,51 | 0,0164 | 0,0083 |
| **E507K-A570T-M747Q** | 4147,97 | 1,12 | 1,45 | 3,54 | 2,88 | 2,55 | 2,73 | 8,02 | 6,18 | 6,29 | 17,61 | 17,55 | 30,08 | 28,47 | 71,53 | 0,40 | 0,0172 | 0,0069 |
| **S515N-M747S** | 4150,20 | 0,63 | 1,47 | 3,05 | 3,11 | 3,67 | 3,52 | 7,61 | 8,94 | 10,97 | 14,94 | 16,85 | 25,24 | 32,00 | 68,00 | 0,47 | 0,0164 | 0,0077 |
| **D732N** | 4328,99 | 0,71 | 1,26 | 2,53 | 2,79 | 3,31 | 2,99 | 7,59 | 7,85 | 9,61 | 14,17 | 17,10 | 30,09 | 29,03 | 70,97 | 0,41 | 0,0164 | 0,0067 |
| **D578Q-D732N** | 4334,18 | 0,70 | 1,34 | 2,20 | 2,01 | 1,60 | 2,49 | 7,08 | 5,92 | 5,87 | 17,12 | 21,69 | 31,98 | 23,35 | 76,65 | 0,30 | 0,0177 | 0,0054 |
| **E708Q-D732N** | 4382,99 | 0,88 | 1,45 | 2,74 | 2,39 | 2,34 | 2,70 | 8,14 | 6,41 | 5,75 | 17,05 | 18,52 | 31,62 | 27,06 | 72,94 | 0,37 | 0,0166 | 0,0062 |
| **E507K-A570G-M747Q** | 4459,80 | 1,33 | 1,27 | 3,71 | 2,92 | 3,00 | 2,58 | 8,65 | 7,26 | 6,59 | 18,74 | 16,32 | 27,65 | 30,71 | 69,29 | 0,44 | 0,0155 | 0,0069 |
| **E507K-D578E** | 4636,97 | 0,87 | 1,44 | 3,40 | 1,96 | 2,36 | 2,45 | 7,27 | 7,04 | 5,08 | 17,35 | 20,90 | 29,89 | 26,78 | 73,22 | 0,37 | 0,0158 | 0,0058 |
| **D578N** | 4677,34 | 1,47 | 1,60 | 2,55 | 2,81 | 4,55 | 3,57 | 7,18 | 7,17 | 5,77 | 14,15 | 21,48 | 27,69 | 30,91 | 69,09 | 0,45 | 0,0148 | 0,0066 |
| **WT** | 4912,92 | 0,71 | 1,37 | 2,51 | 2,53 | 3,01 | 3,82 | 6,36 | 6,05 | 5,92 | 14,82 | 22,83 | 30,09 | 26,34 | 73,66 | 0,36 | 0,0150 | 0,0054 |
| **M747Q** | 4985,04 | 1,18 | 1,38 | 2,49 | 2,40 | 4,31 | 3,83 | 7,18 | 7,36 | 4,82 | 15,22 | 21,98 | 27,85 | 30,13 | 69,87 | 0,43 | 0,0140 | 0,0060 |
| **E708D** | 5000,88 | 0,58 | 1,34 | 2,06 | 1,83 | 1,86 | 2,26 | 7,18 | 5,43 | 5,37 | 16,32 | 20,35 | 35,42 | 22,54 | 77,46 | 0,29 | 0,0155 | 0,0045 |
| **D578K** | 5140,39 | 0,68 | 1,42 | 1,65 | 2,85 | 4,11 | 4,37 | 5,77 | 5,01 | 7,80 | 14,89 | 20,87 | 30,58 | 25,86 | 74,14 | 0,35 | 0,0144 | 0,0050 |
| **E708L** | 5164,52 | 0,64 | 1,33 | 2,28 | 2,01 | 1,88 | 2,67 | 7,07 | 5,32 | 6,18 | 16,69 | 19,47 | 34,46 | 23,21 | 76,79 | 0,30 | 0,0149 | 0,0045 |
| **V586N-M747T** | 5192,41 | 0,51 | 1,18 | 1,74 | 3,40 | 2,08 | 2,78 | 5,56 | 3,98 | 10,36 | 15,79 | 22,22 | 30,39 | 21,24 | 78,76 | 0,27 | 0,0152 | 0,0041 |
| **E507K** | 5205,19 | 1,00 | 1,41 | 2,82 | 2,54 | 4,72 | 3,78 | 6,71 | 7,26 | 5,47 | 15,90 | 20,50 | 27,91 | 30,22 | 69,78 | 0,43 | 0,0134 | 0,0058 |
| **E626K** | 5228,77 | 0,60 | 1,42 | 2,00 | 1,95 | 1,75 | 2,91 | 6,52 | 5,72 | 5,13 | 15,81 | 21,66 | 34,51 | 22,88 | 77,12 | 0,30 | 0,0147 | 0,0044 |
| **D578W** | 5440,88 | 0,97 | 1,53 | 1,88 | 2,79 | 4,22 | 3,41 | 7,10 | 7,20 | 6,05 | 15,32 | 20,01 | 29,51 | 29,10 | 70,90 | 0,41 | 0,0130 | 0,0053 |
| **E507Q** | 5633,62 | 0,89 | 1,60 | 2,48 | 1,89 | 4,39 | 4,04 | 6,18 | 6,70 | 4,78 | 15,64 | 21,98 | 29,43 | 28,17 | 71,83 | 0,39 | 0,0128 | 0,0050 |
| **D578S-R728Q-M747Q** | 5946,68 | 0,44 | 1,46 | 1,39 | 1,54 | 1,16 | 2,49 | 4,76 | 2,76 | 7,54 | 18,19 | 19,55 | 38,74 | 15,99 | 84,01 | 0,19 | 0,0141 | 0,0027 |
| **S515F** | 6114,36 | 0,76 | 1,45 | 1,62 | 2,08 | 3,90 | 3,63 | 5,10 | 5,02 | 5,69 | 21,61 | 22,94 | 26,20 | 23,56 | 76,44 | 0,31 | 0,0125 | 0,0039 |
| **D578S-R728Q-M747V** | 6346,43 | 0,30 | 1,25 | 1,51 | 1,73 | 0,96 | 2,72 | 2,82 | 2,85 | 5,67 | 19,85 | 21,84 | 38,49 | 14,15 | 85,85 | 0,16 | 0,0135 | 0,0022 |
| **E507D** | 6644,88 | 0,77 | 1,59 | 1,23 | 2,57 | 2,05 | 3,32 | 5,00 | 3,68 | 7,31 | 17,30 | 23,51 | 31,66 | 20,21 | 79,79 | 0,25 | 0,0120 | 0,0030 |
| **E507Q-D578S-R728Q** | 6889,79 | 0,34 | 1,57 | 1,26 | 1,66 | 0,87 | 2,82 | 2,70 | 2,63 | 7,24 | 17,04 | 22,36 | 39,50 | 13,85 | 86,15 | 0,16 | 0,0125 | 0,0020 |
| **I707L** | 7441,63 | 0,66 | 1,55 | 3,31 | 2,64 | 2,40 | 3,47 | 4,96 | 4,18 | 6,67 | 21,69 | 17,12 | 31,36 | 23,16 | 76,84 | 0,30 | 0,0103 | 0,0031 |
| **M747E** | 7543,12 | 0,25 | 1,56 | 1,17 | 1,91 | 1,77 | 3,99 | 4,16 | 2,72 | 4,29 | 17,36 | 27,20 | 33,64 | 17,52 | 82,48 | 0,21 | 0,0109 | 0,0023 |
| **S515N** | 7742,98 | 0,34 | 1,58 | 1,29 | 2,04 | 2,35 | 4,49 | 3,96 | 2,94 | 5,11 | 17,22 | 26,24 | 32,45 | 18,98 | 81,02 | 0,23 | 0,0105 | 0,0025 |
| **R573K-D578N** | 7828,28 | 0,32 | 1,93 | 1,33 | 1,10 | 1,25 | 3,60 | 5,77 | 4,49 | 3,58 | 20,17 | 15,16 | 41,30 | 19,79 | 80,21 | 0,25 | 0,0102 | 0,0025 |
| **F667Y** | 8433,00 | 0,41 | 1,81 | 1,40 | 2,78 | 1,36 | 4,54 | 6,32 | 4,28 | 8,49 | 21,80 | 21,26 | 25,56 | 22,88 | 77,12 | 0,30 | 0,0091 | 0,0027 |
| **H639F** | 8614,00 | 0,08 | 0,82 | 0,44 | 2,21 | 0,66 | 4,09 | 1,19 | 0,83 | 3,84 | 18,49 | 17,44 | 49,90 | 10,32 | 89,68 | 0,12 | 0,0104 | 0,0012 |
| **R660S** | 10996,46 | 0,56 | 1,98 | 3,39 | 3,73 | 2,42 | 3,84 | 4,21 | 3,49 | 11,04 | 27,87 | 16,62 | 20,86 | 23,62 | 76,38 | 0,31 | 0,0069 | 0,0021 |
| **E507K-R536K-R660V** | 13051,67 | 0,26 | 1,31 | 6,85 | 1,67 | 1,38 | 2,16 | 3,00 | 6,01 | 7,70 | 49,40 | 13,42 | 6,84 | 22,65 | 77,35 | 0,29 | 0,0059 | 0,0017 |
| **E507R-R573K-E708Q** | 13458,91 | 0,32 | 2,41 | 2,23 | 1,94 | 1,44 | 4,75 | 5,10 | 3,80 | 3,26 | 28,47 | 11,50 | 34,78 | 21,98 | 78,02 | 0,28 | 0,0058 | 0,0016 |
| **F667M** | 13759,44 | 0,39 | 3,61 | 0,69 | 3,20 | 1,53 | 6,46 | 2,74 | 1,80 | 5,56 | 26,12 | 18,14 | 29,75 | 20,43 | 79,57 | 0,26 | 0,0058 | 0,0015 |
| **H639A** | 14470,52 | 0,12 | 1,53 | 1,91 | 2,74 | 0,93 | 6,13 | 1,55 | 1,41 | 4,85 | 23,31 | 15,46 | 40,06 | 16,32 | 83,68 | 0,19 | 0,0058 | 0,0011 |
| **S515D** | 14600,83 | 0,25 | 2,32 | 2,60 | 2,59 | 2,40 | 6,21 | 2,40 | 1,54 | 4,32 | 28,98 | 13,26 | 33,12 | 20,31 | 79,69 | 0,25 | 0,0055 | 0,0014 |
| **M747D** | 15244,39 | 0,08 | 2,47 | 0,86 | 2,68 | 1,49 | 7,68 | 2,04 | 2,16 | 2,57 | 27,64 | 19,59 | 30,75 | 19,45 | 80,55 | 0,24 | 0,0053 | 0,0013 |
| **R660V** | 15402,44 | 0,54 | 1,99 | 5,04 | 2,95 | 1,13 | 5,44 | 4,03 | 4,14 | 7,02 | 35,50 | 15,62 | 16,60 | 25,27 | 74,73 | 0,34 | 0,0049 | 0,0016 |
| **F667I-R728N-M747S** | 22506,31 | 0,53 | 1,71 | 1,84 | 3,11 | 2,23 | 6,99 | 4,24 | 4,57 | 12,42 | 26,45 | 17,24 | 18,69 | 25,20 | 74,80 | 0,34 | 0,0033 | 0,0011 |
| **S515D-K540M** | 28816,50 | 0,38 | 1,07 | 1,09 | 5,51 | 7,68 | 11,36 | 2,86 | 5,08 | 15,15 | 7,50 | 20,04 | 22,28 | 35,02 | 64,98 | 0,54 | 0,0023 | 0,0012 |
| **F667A** | 42014,26 | 0,54 | 4,38 | 2,73 | 6,93 | 2,51 | 10,18 | 3,76 | 2,10 | 14,36 | 39,10 | 6,82 | 6,58 | 33,14 | 66,86 | 0,50 | 0,0016 | 0,0008 |

**Table S3.** Polymerase synthesis fidelity assessments of 47 Taq pol variants. The enzymes are stratified from top to bottom in increasing fidelity. Enzymes that have demonstrated sufficient RT activity are labeled in red. Fidelity is given as the error rate (1 error/n nucleotides). The relative frequency of individual variants of misincorporations for each enzyme is given (the total frequency of all errors is taken as 100%).TV, transversions; TS, transitions. %TS (%TV) – the total frequency of transitions (transversions). TV/TS – transversions/transitions ratio.
