## Supplementary material for "Enhancing the Reverse Transcriptase Function in Taq Polymerase via AI-driven Multiparametric Rational Design": Table S4

| **Position** | **Сount** |  |  |  |  |  |  |  |  |  |  |  |  |  |  |  |  |  |  |  |  |  |  |  |  |  |  |  |  |  |  |  |  |  |  |  |  |  |  |
| --- | --- | --- | --- | --- | --- | --- | --- | --- | --- | --- | --- | --- | --- | --- | --- | --- | --- | --- | --- | --- | --- | --- | --- | --- | --- | --- | --- | --- | --- | --- | --- | --- | --- | --- | --- | --- | --- | --- | --- |
| **E742** | 33007 | **EQ** | 6730 | **EM** | 3965 | **EH** | 3299 | **EN** | 2957 | **EY** | 2863 | **EL** | 2174 | **ES** | 2043 | **EA** | 1731 | **EW** | 1288 | **EF** | 1259 | **EK** | 1207 | **ET** | 1154 | **EG** | 877 | **ER** | 683 | **ED** | 312 | **EP** | 239 | **EI** | 153 | **EV** | 73 |  |  |
| **D578** | 24424 | **DN** | 2808 | **DQ** | 2721 | **DH** | 2418 | **DW** | 2231 | **DR** | 2217 | **DK** | 1904 | **DG** | 1608 | **DY** | 1535 | **DM** | 1472 | **DS** | 1470 | **DA** | 1162 | **DE** | 858 | **DT** | 600 | **DP** | 414 | **DF** | 385 | **DL** | 312 | **DV** | 298 | **DI** | 11 |  |  |
| **E507** | 22257 | **EK** | 7197 | **ER** | 4445 | **ET** | 3144 | **EH** | 2515 | **EQ** | 1893 | **EA** | 693 | **ED** | 556 | **EW** | 535 | **EY** | 479 | **EL** | 186 | **EF** | 178 | **EV** | 172 | **EN** | 142 | **EP** | 81 | **EM** | 31 | **EI** | 10 |  |  |  |  |  |  |
| **I707** | 18979 | **IR** | 5539 | **IK** | 3483 | **IM** | 1918 | **IQ** | 1366 | **IH** | 1230 | **IL** | 1101 | **IW** | 995 | **IV** | 782 | **IA** | 539 | **IG** | 498 | **IT** | 413 | **IF** | 410 | **IY** | 405 | **IS** | 148 | **IN** | 131 | **IE** | 19 | **IP** | 2 |  |  |  |  |
| **A570** | 17833 | **AK** | 4735 | **AR** | 4011 | **AN** | 1319 | **AM** | 933 | **AQ** | 1765 | **AH** | 1216 | **AW** | 705 | **AG** | 550 | **AT** | 498 | **AY** | 475 | **AV** | 425 | **AP** | 364 | **AS** | 321 | **AL** | 265 | **AI** | 129 | **AF** | 115 | **AD** | 6 | **AE** | 1 |  |  |
| **A743** | 15150 | **AQ** | 2143 | **AM** | 1570 | **AR** | 1376 | **AY** | 1374 | **AH** | 1364 | **AN** | 1305 | **AK** | 992 | **AF** | 893 | **AT** | 823 | **AS** | 775 | **AL** | 702 | **AG** | 586 | **AW** | 549 | **AV** | 405 | **AP** | 140 | **AI** | 131 | **AE** | 12 | **AD** | 10 |  |  |
| **M747** | 13418 | **MK** | 3617 | **MR** | 2659 | **MQ** | 2509 | **MH** | 1138 | **MY** | 511 | **MT** | 477 | **MA** | 445 | **ML** | 401 | **MF** | 382 | **MV** | 299 | **MS** | 292 | **MN** | 202 | **MI** | 184 | **MG** | 175 | **MW** | 120 | **MP** | 7 |  |  |  |  |  |  |
| **E708** | 10205 | **EQ** | 3146 | **EK** | 1428 | **EA** | 1428 | **ER** | 1105 | **EH** | 1052 | **EM** | 508 | **EG** | 507 | **ED** | 318 | **EW** | 213 | **ES** | 206 | **EP** | 115 | **EL** | 90 | **EN** | 44 | **EY** | 22 | **EF** | 17 | **ET** | 6 |  |  |  |  |  |  |
| **V586** | 8227 | **VR** | 1772 | **VK** | 1535 | **VQ** | 747 | **VM** | 656 | **VI** | 562 | **VL** | 552 | **VH** | 442 | **VN** | 414 | **VA** | 263 | **VT** | 257 | **VW** | 257 | **VY** | 200 | **VF** | 172 | **VG** | 164 | **VP** | 149 | **VS** | 85 |  |  |  |  |  |  |
| **S515** | 5914 | **SN** | 1571 | **SK** | 958 | **SQ** | 915 | **SR** | 663 | **SH** | 516 | **ST** | 465 | **SA** | 240 | **SD** | 164 | **SP** | 156 | **SG** | 118 | **SE** | 86 | **SM** | 37 | **SY** | 16 | **SV** | 9 |  |  |  |  |  |  |  |  |  |  |
| **I614** | 5202 | **IM** | 2237 | **IV** | 769 | **IL** | 688 | **IF** | 241 | **IQ** | 405 | **IW** | 195 | **IK** | 192 | **IT** | 174 | **IR** | 125 | **IN** | 59 | **IE** | 50 | **IA** | 36 | **IC** | 15 | **IS** | 11 | **IY** | 3 | **IH** | 1 | **IG** | 1 |  |  |  |  |
| **Q754** | 2734 | **QK** | 576 | **QR** | 406 | **QH** | 377 | **QM** | 344 | **QE** | 203 | **QN** | 165 | **QP** | 138 | **QS** | 115 | **QW** | 115 | **QA** | 88 | **QY** | 79 | **QL** | 56 | **QG** | 38 | **QT** | 15 | **QC** | 7 | **QV** | 5 | **QD** | 4 | **QF** | 3 |  |  |
| **E626** | 1923 | **EP** | 951 | **EQ** | 571 | **EA** | 235 | **ED** | 64 | **EK** | 43 | **EH** | 21 | **EW** | 20 | **ES** | 9 | **EV** | 5 | **ER** | 4 |  |  |  |  |  |  |  |  |  |  |  |  |  |  |  |  |  |  |
| **V783** | 1809 | **VI** | 614 | **VM** | 189 | **VL** | 168 | **VQ** | 134 | **VW** | 109 | **VT** | 95 | **VY** | 82 | **VR** | 65 | **VH** | 64 | **VN** | 61 | **VF** | 59 | **VA** | 53 | **VK** | 48 | **VC** | 38 | **VS** | 16 | **VE** | 5 | **VD** | 4 | **VG** | 3 | **VP** | 2 |
| **R573** | 1672 | **RK** | 674 | **RH** | 541 | **RW** | 190 | **RP** | 187 | **RQ** | 30 | **RC** | 20 | **RD** | 18 | **RG** | 6 | **RL** | 4 | **RS** | 2 |  |  |  |  |  |  |  |  |  |  |  |  |  |  |  |  |  |  |
| **N483** | 912 | **NK** | 199 | **NT** | 173 | **NH** | 139 | **NR** | 122 | **NQ** | 101 | **NS** | 66 | **NM** | 43 | **NA** | 20 | **NG** | 19 | **NY** | 10 | **NW** | 9 | **NL** | 5 | **NV** | 5 | **NE** | 1 |  |  |  |  |  |  |  |  |  |  |
| **F667** | 624 | **FY** | 624 |  |  |  |  |  |  |  |  |  |  |  |  |  |  |  |  |  |  |  |  |  |  |  |  |  |  |  |  |  |  |  |  |  |  |  |  |
| **K540** | 172 | **KR** | 172 |  |  |  |  |  |  |  |  |  |  |  |  |  |  |  |  |  |  |  |  |  |  |  |  |  |  |  |  |  |  |  |  |  |  |  |  |
| **R728** | 106 | **RK** | 103 | **RH** | 3 |  |  |  |  |  |  |  |  |  |  |  |  |  |  |  |  |  |  |  |  |  |  |  |  |  |  |  |  |  |  |  |  |  |  |
| **R746** | 92 | **RK** | 92 |  |  |  |  |  |  |  |  |  |  |  |  |  |  |  |  |  |  |  |  |  |  |  |  |  |  |  |  |  |  |  |  |  |  |  |  |
| **D732** | 87 | **DN** | 14 | **DG** | 13 | **DH** | 11 | **DA** | 9 | **DQ** | 9 | **DT** | 7 | **DS** | 6 | **DM** | 6 | **DK** | 3 | **DR** | 2 | **DY** | 2 | **DW** | 1 | **DP** | 1 | **DC** | 1 | **DV** | 1 | **DE** | 1 |  |  |  |  |  |  |
| **H784** | 54 | **HQ** | 34 | **HR** | 6 | **HW** | 6 | **HY** | 4 | **HK** | 2 | **HN** | 2 |  |  |  |  |  |  |  |  |  |  |  |  |  |  |  |  |  |  |  |  |  |  |  |  |  |  |
| **H639** | 12 | **HQ** | 5 | **HM** | 5 | **HR** | 1 | **HY** | 1 |  |  |  |  |  |  |  |  |  |  |  |  |  |  |  |  |  |  |  |  |  |  |  |  |  |  |  |  |  |  |
| **R659** | 3 | **RK** | 3 |  |  |  |  |  |  |  |  |  |  |  |  |  |  |  |  |  |  |  |  |  |  |  |  |  |  |  |  |  |  |  |  |  |  |  |  |
| **E159** | 2 | **EK** | 1 | **ER** | 1 |  |  |  |  |  |  |  |  |  |  |  |  |  |  |  |  |  |  |  |  |  |  |  |  |  |  |  |  |  |  |  |  |  |  |
| **E189** | 2 | **ER** | 1 | **EK** | 1 |  |  |  |  |  |  |  |  |  |  |  |  |  |  |  |  |  |  |  |  |  |  |  |  |  |  |  |  |  |  |  |  |  |  |
| **A141** | 1 | **AR** | 1 |  |  |  |  |  |  |  |  |  |  |  |  |  |  |  |  |  |  |  |  |  |  |  |  |  |  |  |  |  |  |  |  |  |  |  |  |
| **A568** | 1 | **AK** | 1 |  |  |  |  |  |  |  |  |  |  |  |  |  |  |  |  |  |  |  |  |  |  |  |  |  |  |  |  |  |  |  |  |  |  |  |  |
| **D551** | 1 | **DK** | 1 |  |  |  |  |  |  |  |  |  |  |  |  |  |  |  |  |  |  |  |  |  |  |  |  |  |  |  |  |  |  |  |  |  |  |  |  |
| **E57** | 1 | **ER** | 1 |  |  |  |  |  |  |  |  |  |  |  |  |  |  |  |  |  |  |  |  |  |  |  |  |  |  |  |  |  |  |  |  |  |  |  |  |
| **E745** | 1 | **EK** | 1 |  |  |  |  |  |  |  |  |  |  |  |  |  |  |  |  |  |  |  |  |  |  |  |  |  |  |  |  |  |  |  |  |  |  |  |  |

**Тable S4.** Positions and types of amino acid substitutions predicted to significantly enhance RTase activity compared to wild-type Taq polymerase. For each substitution, the amino acid to be replaced, the replacing amino acid, and the number of predicted enzymes with enhanced RTase activity carrying such substitution (including variants with single, double, and triple substitutions) are indicated. The criterion of enhanced RTase activity was taken dCt < 2 between the enzyme tested and the control p66 RTase at 90 b. and 116 b. cDNA synthesis and further real-time PCR amplification by the wild-type Taq polymerase (see **Methods** section). This characteristic was predicted for > 60,000 candidates. They carried 296 amino acid substitutions of 174 types in 31 different positions (1 to 19 substitutions per position).
