## Supplementary figures and images for "Enhancing the Reverse Transcriptase Function in Taq Polymerase via AI-driven Multiparametric Rational Design"

### Figure S1

#### Slide 1
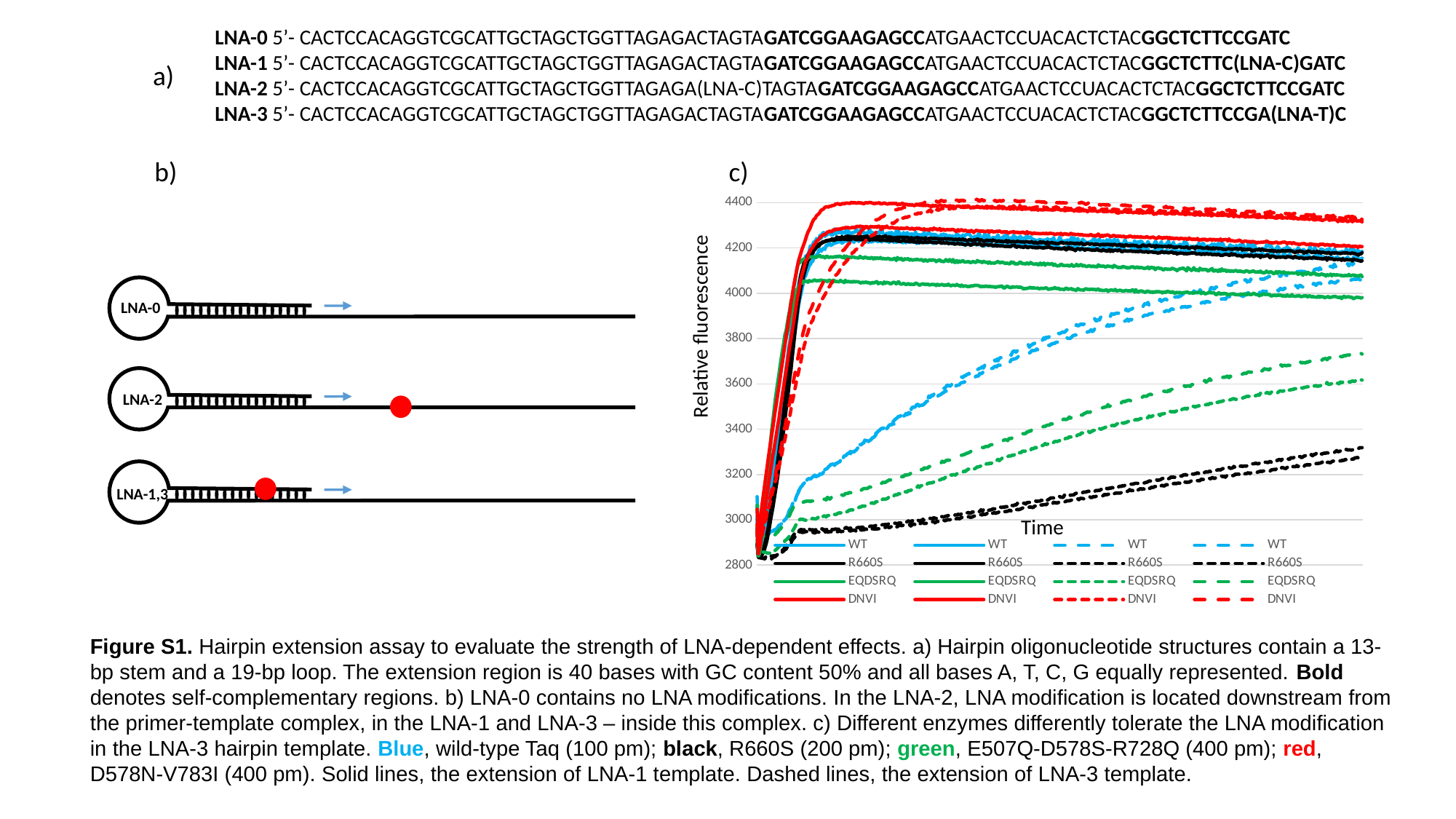
