## Supplementary material for "Enhancing the Reverse Transcriptase Function in Taq Polymerase via AI-driven Multiparametric Rational Design": Figure S2

### Slide 1
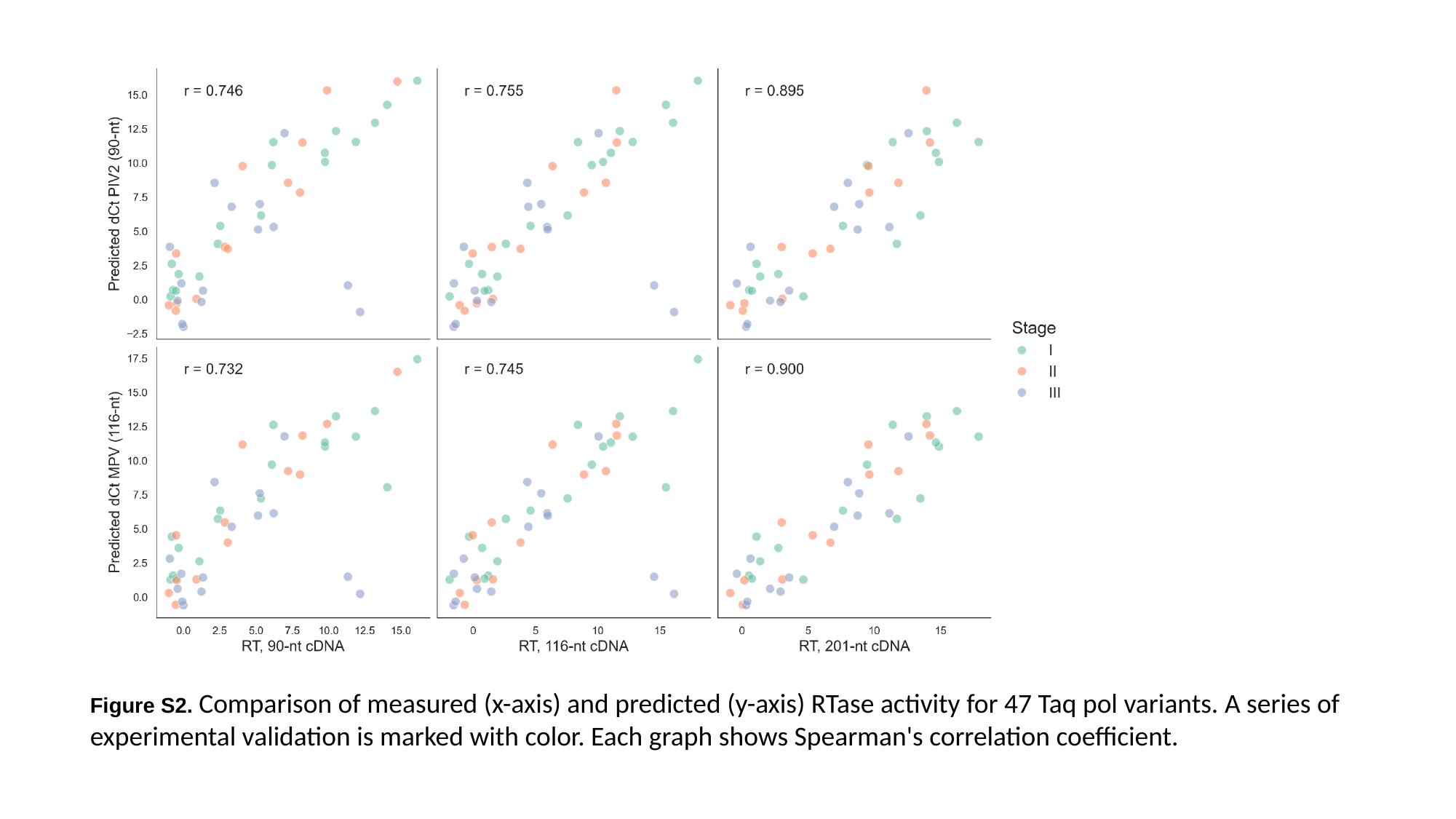

Figure S2. Comparison of measured (x-axis) and predicted (y-axis) RTase activity for 47 Taq pol variants. A series of experimental validation is marked with color. Each graph shows Spearman's correlation coefficient.
