## Supplementary material for "Enhancing the Reverse Transcriptase Function in Taq Polymerase via AI-driven Multiparametric Rational Design": Figure S3

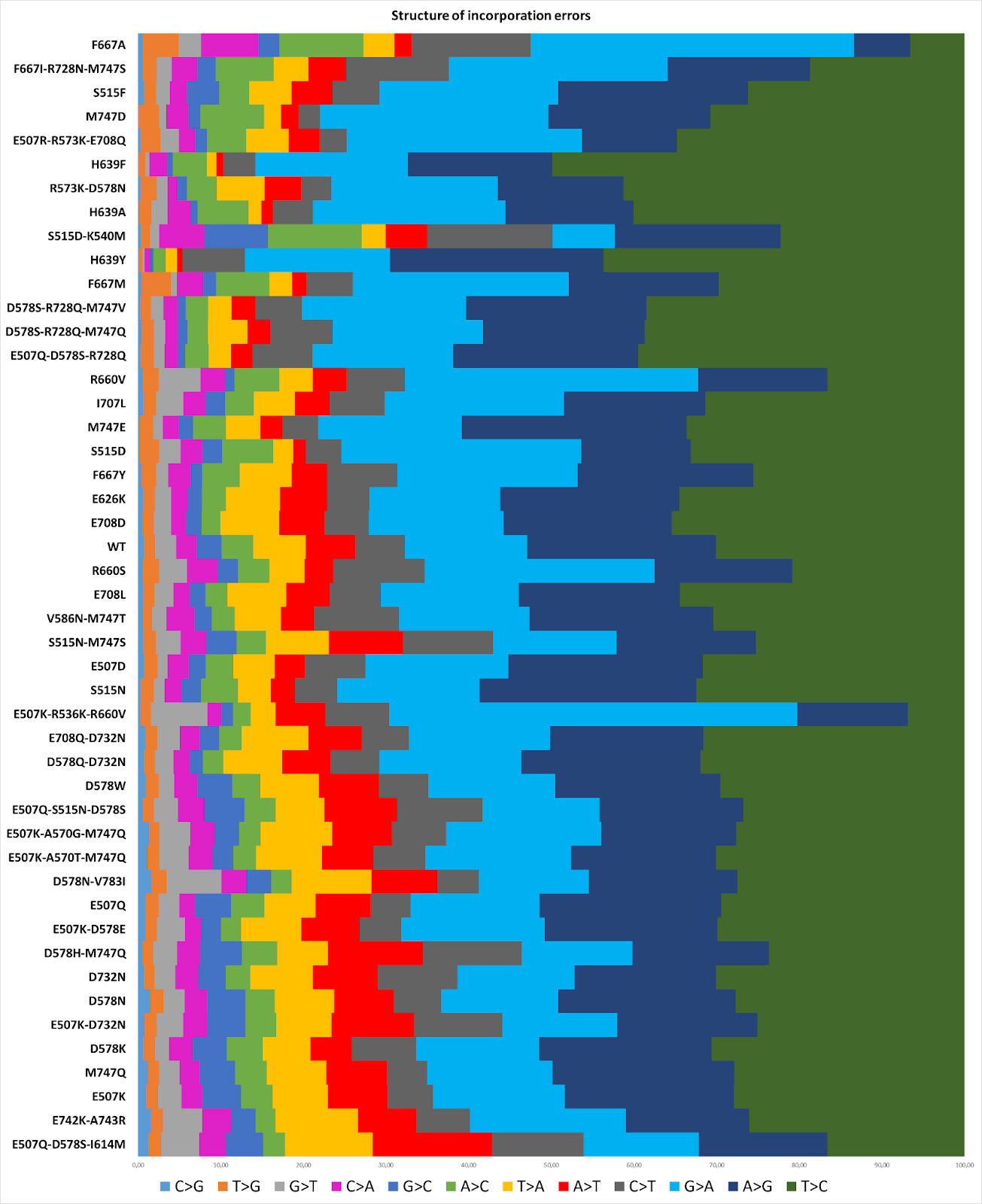

**Figure S3.** Structure of incorporation errors made by different Taq pol variants. The enzymes are stratified from top to bottom by an increasing RTase activity.
