## Supplementary material for "Enhancing the Reverse Transcriptase Function in Taq Polymerase via AI-driven Multiparametric Rational Design": Figure S4

### Slide 1
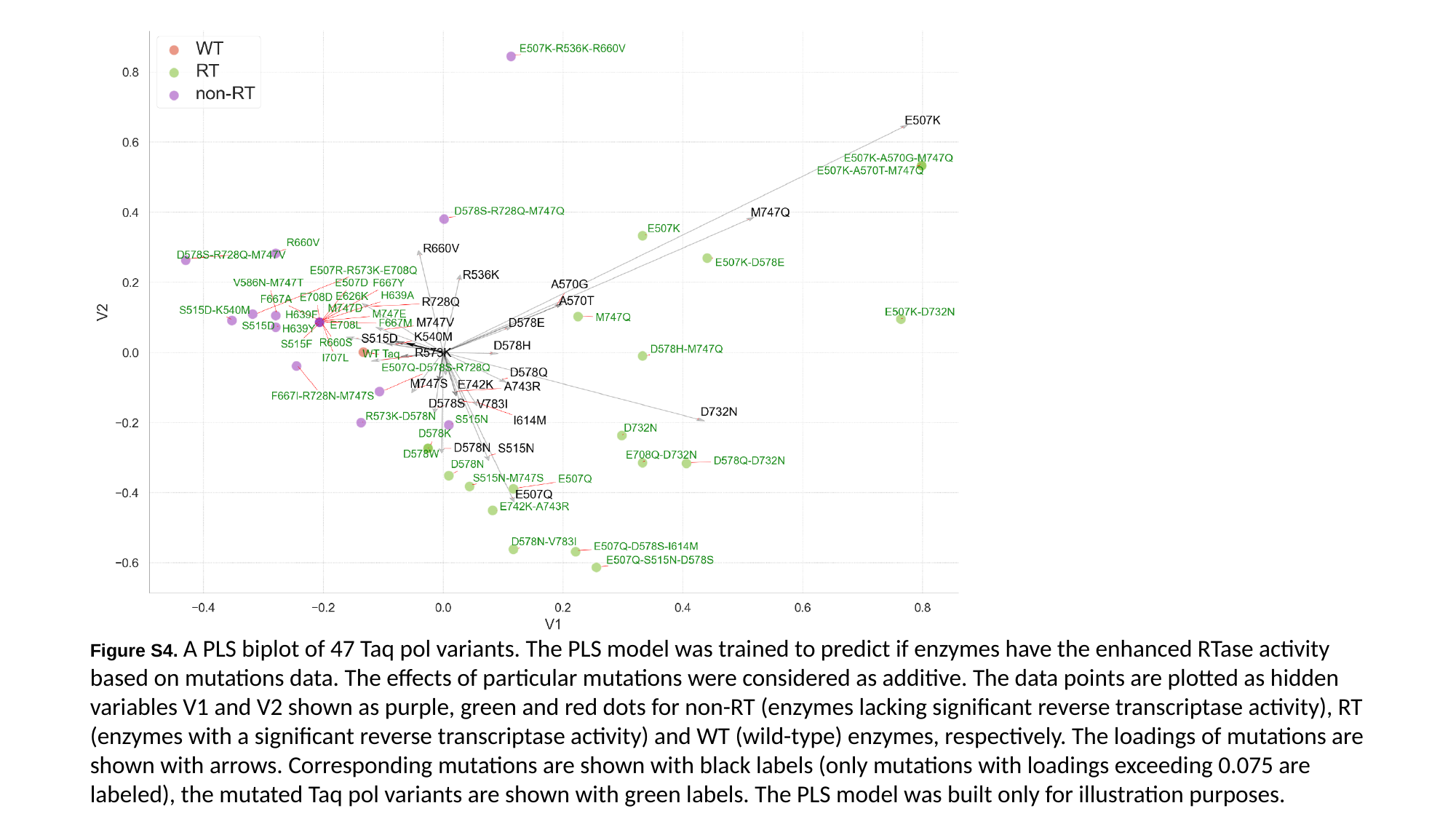

Figure S4. A PLS biplot of 47 Taq pol variants. The PLS model was trained to predict if enzymes have the enhanced RTase activity based on mutations data. The effects of particular mutations were considered as additive. The data points are plotted as hidden variables V1 and V2 shown as purple, green and red dots for non-RT (enzymes lacking significant reverse transcriptase activity), RT (enzymes with a significant reverse transcriptase activity) and WT (wild-type) enzymes, respectively. The loadings of mutations are shown with arrows. Corresponding mutations are shown with black labels (only mutations with loadings exceeding 0.075 are labeled), the mutated Taq pol variants are shown with green labels. The PLS model was built only for illustration purposes.
